# Passivating blunt-ended helices to control monodispersity and multi-subunit assembly of DNA origami structures

**DOI:** 10.1101/2023.09.09.557003

**Authors:** Jonathan F. Berengut, Willi R. Berg, Felix J. Rizzuto, Lawrence K. Lee

## Abstract

DNA origami facilitates the synthesis of bespoke nanoscale structures suitable for a wide range of applications. Effective design requires prevention of uncontrolled aggregation, while still permitting directed assembly of multi-subunit superstructures. Uncontrolled aggregation can be caused by base-stacking interactions between arrays of blunt-ended helices on different structures, which are routinely passivated by incorporating disordered regions as either scaffold loops or poly-nucleotide brushes (usually poly-thymine) at the end of DNA helices. Such disordered regions are ubiquitous in DNA origami structures yet their exact design requirements in different chemical environments are ill defined. In this study, we systematically examine the use of scaffold loops and poly-nucleotide brushes for passivation and for controlling multi-subunit assembly. We assess the dependence of length and sequence for preventing aggregation amidst a titration of MgCl_2_ concentrations and the suitability of each strategy for enabling controlled multi-subunit assembly. We then introduce a novel strategy where double-stranded DNA helices run orthogonal to arrays of blunt-ended DNA helices forming a steric shield that prevents base stacking. The results define the limitations of each method and important design considerations for achieving monodispersity. For example, poly-thymine brushes are most effective for achieving monodispersity in the broadest conditions whereas scaffold loops can facilitate directed multi-subunit assembly. Finally, orthogonal DNA helices remove the need for disordered regions altogether, prevent aggregation in a broad range of MgCl_2_ concentrations and facilitate directed multi-subunit assembly. This study expands the design tools available and enables a more informed approach for achieving control of monodispersity and multi-subunit assembly in DNA origami structures.

## Introduction

DNA nanotechnology exploits the well-known structure and base-pairing properties of DNA to create self-assembling nanoscale structures with defined, bespoke geometries.^[1]^ DNA origami^[2]^ is a popular method where long single-stranded DNA (ssDNA) scaffolds are mixed with a molar excess of shorter ssDNA staples, which anneal according to Watson-Crick base-pairing rules and ‘fold’ into predetermined structures. Each origami structure comprises spatially organized double-stranded DNA (dsDNA) helices, usually arranged into a flexible two-dimensional sheet, or a more rigid three-dimensional lattice or mesh^[3, 4]^ **(Figure 1A)**. DNA origami is particularly useful because it facilitates the design and high-yield synthesis of complex, nanoscale 3D structures. Moreover, DNA origami structures have site-specific addressability, enabling precise control over the location of nanoparticles and biomolecules in 3D, making them suitable for a myriad of applications in biosensing,^[5]^ biophysics,^[6]^ and molecular medicine.^[7]^

**Figure 1.**
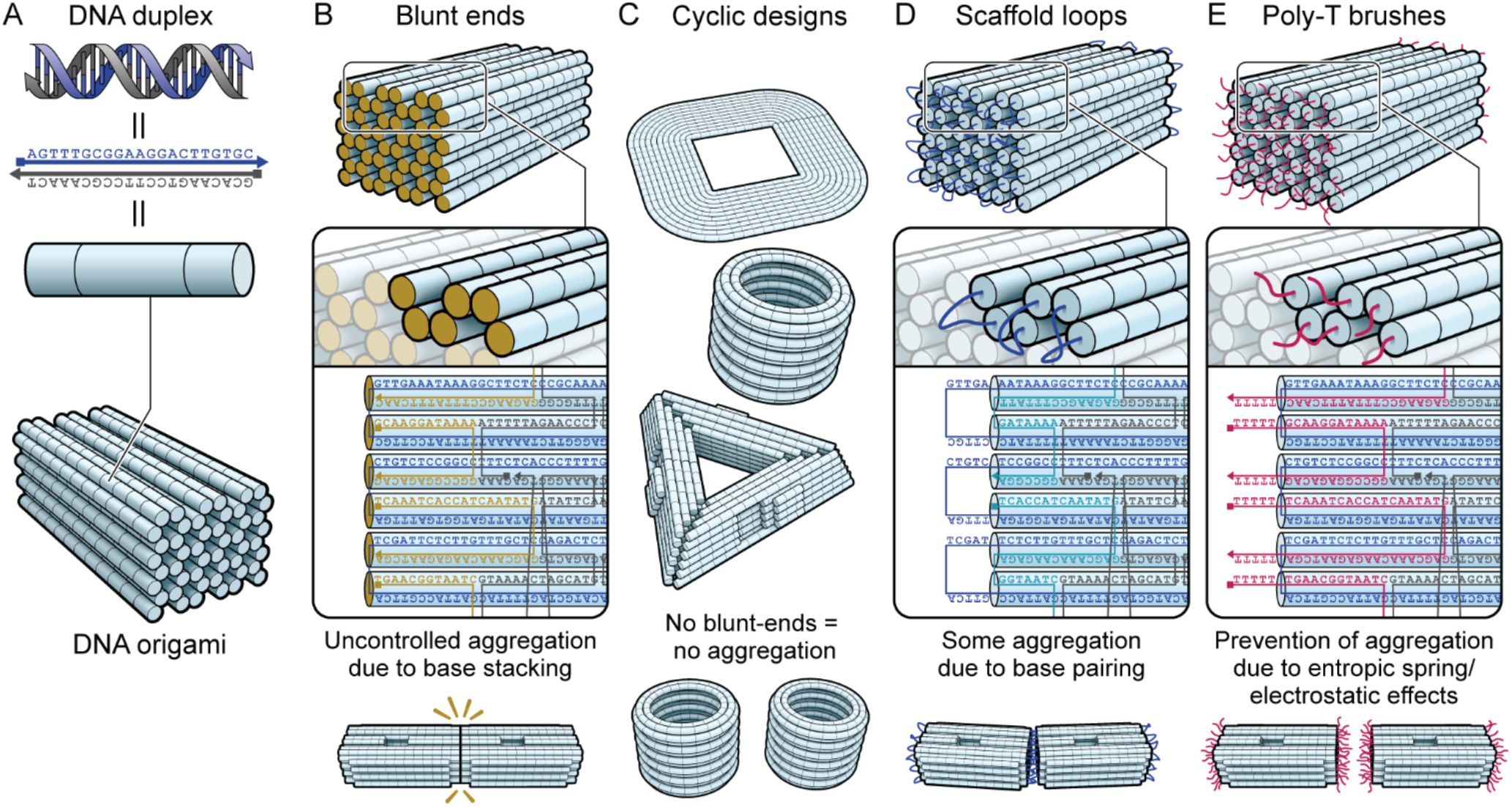
Principles and schematics of aggregation prevention in DNA origami design. A) Presentation of various visual representations of DNA and DNA origami. B) Blunt-ends occur at the termini of double-stranded DNA, which are often present at the ends of DNA origami structures. These blunt-ends, when occurring in large numbers, tend to cause aggregation due to base-stacking with other origami. C) Examples of DNA origami designs that minimise or eliminate blunt ends but may not be suitable for all applications. Designs top-bottom are from Han *et al*.^[12]^, Wickham *et al*.^[11]^ and Sigl *et al*.^[14]^, respectively. D) By truncating (or omitting) staple strands at the ends of the origami, single-stranded scaffold loops are exposed. These are moderately successful in preventing aggregation, but can cause aggregation due to inadvertent base-pairing. E) By extending the staples at the ends of the origami with polymers of identical nucleotides (for example poly-thymine), aggregation is effectively prevented.

Regulating interactions between DNA origami structures is important to prevent uncontrolled, non-specific aggregation and if required, to enable the controlled self-assembly of multi-subunit structures. Uncontrolled aggregation can result from poorly folded DNA origami subunits but can also occur in well folded structures, most notably via base stacking interactions. In latticed designs the parallel bundles of dsDNA helices present multiple exposed nucleobases at their terminal ends.^[8]^ Individually, these ‘blunt ends’ have a propensity to form weak stacking interactions with other exposed DNA bases. In latticed structures, many simultaneous intermolecular base-stacking interactions yield stable interactions between DNA origami structures, leading to uncontrolled aggregation **(Figure 1B)**.

A reduction in base-stacking mediated aggregation in the presence of the denaturing agent formamide has been reported.^[9]^ However, chemical conditions that weaken unwanted inter-subunit base-stacking interactions such as lowering salt concentrations, increasing temperature or reducing DNA concentration, also act to prevent proper assembly of target DNA structures. Ionic screening with the addition of Mg^2+^ or Na^+^ is required for efficient DNA hybridisation, for example.^[3, 10]^ Consequently, methods that focus on DNA origami design to control base-stacking interactions are far more common.

One design-based approach to control aggregation is to construct DNA origami structures that by design do not display blunt ends. These can take the form of circular structures,^[11, 12]^ or frame-like structures with mitred or rounded corners^[13, 14]^ **(Figure 1C)**. These designs require that each continuous DNA double-helix follows a looped path, a constraint that may make this approach unsuitable for some applications.

The most common method of reducing base stacking-mediated aggregation involves the creation of disordered regions of ssDNA at the ends of DNA helices to passivate blunt-ends.^[15, 16]^ The strands in these disordered regions sterically shielding ordered arrays of blunt-ends and act as entropic springs, thus preventing the formation of intermolecular base-stacking interactions.

Disordered regions can comprise unpaired scaffold strand, which can be achieved by truncating or omitting staple strands at the ends of DNA helices^[17]^ **(Figure 1D)**. These scaffold loops are usually ~10 nucleotides (nt) in length and have a sequence determined by the scaffold strand, which is generally a derivative of the M13 phage genome. The sequences of scaffold loops are thus effectively random and include a combination of all four nucleotides (A, G, C and T). Consequently, while base stacking interactions are likely minimized, stable intermolecular base-pairing interactions can occur, thus scaffold loops can potentially cause undesired aggregation rather than prevent it .^[16]^

An alternative approach that similarly produces disordered ssDNA regions at the end of DNA helices is increasing the length of staple strands at the ends of the structure, so that they extend beyond where they hybridize to the scaffold strand in the origami structure. These staple extensions can return to the double-stranded core of the structure, forming single-stranded staple loops,^[18]^ or can simply terminate outside of the structure, forming single-stranded brushes^[16]^ **(Figure 1E)**. The advantage of this strategy over the scaffold loop strategy is that the length and sequence of the extensions can be easily controlled to minimize base-pairing as well as base stacking-mediated aggregation.

Staple extensions almost invariably consist of poly-thymine (poly-T) sequences.^[19-22]^ Uniformity in base composition prevents complementary base pairing and thymine is the least likely to form secondary structures that lead to aggregation. This is because pyrimidine bases (thymine & cytosine) have a smaller surface area of aromatic rings than purines (adenine & guanine) and thus form weaker stacking interactions. Moreover, thymine bases form weaker Watson-Crick base-pairing interactions than cytosine bases. Most published works that have employed this strategy have used poly-T brushes or loops with lengths between 4-10 nt. Despite rapid advances in the applications of DNA origami, the limitations of poly-T brushes and scaffold loops and their precise requirements to achieve effective passivation in different chemical conditions remains ill-defined.

In this study we present a systematic comparison of different design strategies for the prevention of aggregation, or passivation, of DNA origami structures. This includes scaffold loops of different lengths and poly-nucleotide brushes of different compositions and lengths. We also introduce a novel method, termed “orthogonal end-capping”, that does not require the inclusion of disordered single-stranded regions and is suitable for non-cyclic designs.

## Results and discussion

### Comparison of scaffold loops and poly-T extensions for preventing aggregation

To compare passivation strategies, we designed several DNA origami structures, all comprising an identical 42-helix bundle layout, arranged in 6 columns and 8 rows of honeycomb lattice as the core structure **(Figure 2A)**. In each variant, the ends of the dsDNA helices are coplanar, providing a large and consistent canvas of DNA helix ends for testing different passivation strategies. To achieve coplanarity of the 48 helices, it was necessary to include four helices on each of two opposite sides of the structure to accommodate scaffold routing, which were shorter in length so as not to contribute to potential stacking interactions at the main interfaces **(Supplementary figure 1)**. We also included an asymmetric hole as a feature to visually determine the orientation of particles in electron micrographs and to assist in particle averaging.

**Figure 2.**
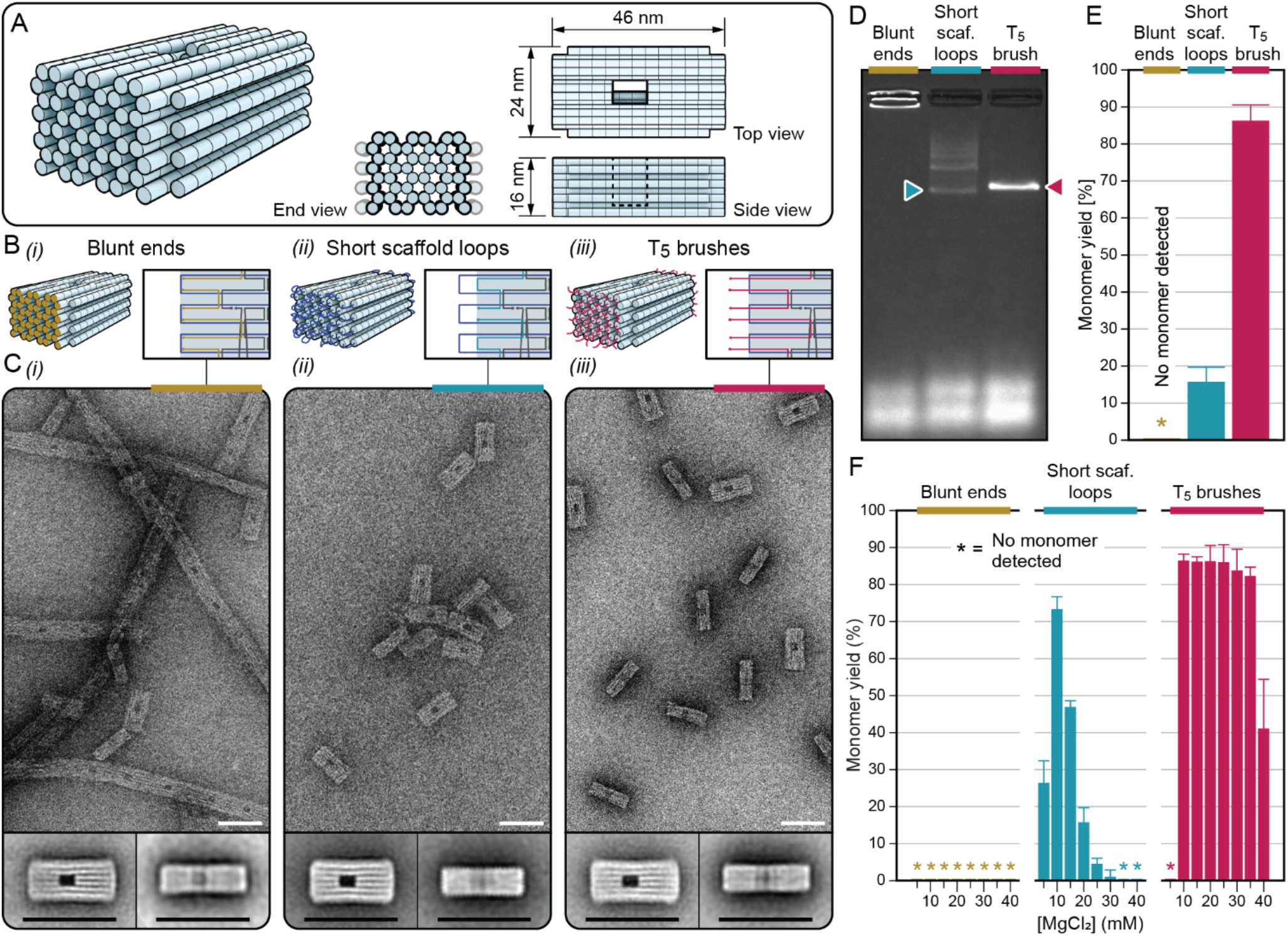
Comparison of three variants of a DNA origami structure with different strategies for preventing aggregation by intermolecular base-stacking. A) Design schematics for the core structure, which was modified to make up each variant. Bi-iii) Schematics of the aggregation prevention strategies presented at the ends of the structures for the blunt-end, short scaffold loop, and T_5_ brush variants respectively. Ci-iii) Exemplary TEM micrographs and 2D particle averages for the three variants. Scale bars are 50 nm. D) An exemplary AGE result used to quantify monomer yield from densitometry of gel bands. E) Monomer yields for the three variants annealed in buffer with 20 mM MgCl_2_. Error bars show standard deviation based on 5 replicates of each variant. F) Monomer yields for the three variants annealed in buffer with MgCl_2_ ranging from 5-40 mM. Error bars show standard deviation based on minimum 3 replicates of each condition.

We initially designed three variants of the structure to compare methods commonly employed to prevent base-stacking interactions. The first was a blunt-ended control, which presented scaffold crossovers coplanar with staple-ends and had no strategies to prevent aggregation from base-stacking **(Figure 2B(i), Supplementary figure 1)**. The second variant employed the scaffold loop method, in which unpaired 10 nt scaffold loops connected neighbouring helices at each end **(Figure 2B(ii), Supplementary figure 2)**. The third variant had 5× poly-T staple extensions (T_5_ brushes) from each helix **(Figure 2B(iii), Supplementary figure 3)**.

Scaffold strands were mixed with a stoichiometric excess of staple strands specific to each DNA origami structure, in buffer with 20 mM MgCl_2_ and thermally annealed to enable folding. The structures were then purified from excess staples and examined via transmission electron microscopy (TEM). All structural variants appeared well-folded, with dimensions corresponding to design and few misfolded structures. The blunt-ended control variant structures formed long polymers with subunits bound at the blunt-ends **(Figure 2C(i), Supplementary figure 4)**. The scaffold-loop variants were mostly monomeric, with occasional aggregates ranging widely in size from 2 to 100’s of subunits **(Figure 2C(ii), Supplementary figure 5)**. Unlike the control variants, aggregates of scaffold loop variants formed disordered clusters rather than well-ordered linear arrays. Such disordered clusters suggest that while scaffold loops prevented base-stacking interactions, they could also enable an additional mode of aggregation, likely via base-pairing of scaffold loops. The T_5_ brush variants were entirely monomeric, with no evidence of polymerization or aggregation **(Figure 2C(iii), Supplementary figure 6)**.

We next analyzed samples via agarose gel electrophoresis (AGE), which provides an indication of the total yield of well-formed monomers in the synthesis reaction. Poorly folded and aggregated structures tend to migrate slower in the gel matrix than well-folded monomeric structures, and hence are well separated, enabling the yield of monomers to be determined with densitometry analysis^[3]^. Densitometry was performed in each lane by first quantifying the integrated value of the total lane intensity between the gel loading well and the large band attributed to excess staples, which tends to migrate much faster due to the smaller molecular weight. The monomer yield was then calculated as the percentage of the well-folded monomeric band intensity against this total lane intensity **(Supplementary figure 7A)**.

The blunt-ended control structure failed to migrate out of the loading well, with the gel lane showing only fast-moving excess staples and bright bands around the well, consistent with the long polymers observed in TEM micrographs, which may be too large to penetrate the gel matrix (**Figure 2D**). The short scaffold loop variant had a monomer yield of 15.8 ± 3.9%, showing laddering in the gel lane above the fastest-moving origami band, indicative of oligomers and larger assemblies (**Figure 2D & E**), as seen in TEM micrographs. The T_5_ brush variant showed one discrete and dominant band between the excess staples and the loading well, with no other discrete band observed, consistent with well-folded origami with very little misfolded or aggregated structures. The monomer yield for this structure was 86.3 ± 4.2%, with the remaining 14% attributed to a slower moving smear in the gel **(Supplementary figure 7B)**, the nature of which was not evident in TEM micrographs. We conclude that under these conditions, only the T_5_ brush variants were able to effectively prevent base-stacking between DNA origami subunits.

### Resistance of passivation to MgCl_2_

The concentration of cations in the folding buffer is well known to influence the folding quality of DNA origami structures as well as their propensity to aggregate. In particular, at low MgCl_2_ concentrations (<5 mM) 3D DNA origami structures do not fold well, whereas at high MgCl_2_ concentrations (>30 mM), they tend to aggregate.^[3]^ To test the resilience of scaffold loops and poly-T extensions to different concentration of MgCl_2_, we annealed the three structural variants described above in MgCl_2_ concentrations ranging from 5 to 40 mM and calculated the monomer yields with densitometry analysis of AGE images **(Supplementary figures 8-10)**.

A comparison of the yields of monomer vs misfolded or aggregated structures is shown in **figure 2F**. The blunt-ended variants did not produce monomers in any conditions, with samples either failing to fold, or aggregating to such an extent that they were unable to migrate through the gel. The scaffold loop variant had a peak yield (73.3 ± 3.4%) of monomers at 10 mM MgCl_2_, a value which decreased rapidly with increasing magnesium. At 5 mM MgCl_2_, the scaffold loop variant produced a long diffuse band which was counted as a 26% monomer yield because it partially overlapped into the gel migration distance region designated to be well-formed monomer. However, this diffuse band was likely dominated by poorly folded DNA origami and the true monomer yield of this sample was likely much lower (Supplementary figure 9). The T_5_ brush variant had a high monomer yield (80-90%) for all MgCl_2_ concentrations between 10 mM and 35 mM. At 40 mM MgCl_2_, the yield of monomer decreased to 40% whereas at 5 mM MgCl_2_ there was no evidence of well-folded monomers in AGE.

To determine the extent to which monomers could be recovered from aggregated structures, the blunt-end and scaffold loop variants were synthesised in buffer with 20 mM MgCl_2_, purified with PEG precipitation^[23]^ and resuspended in buffer with concentrations of MgCl_2_ between 2 mM and 20 mM **(Supplementary figure 11)**. In 2 mM resuspension buffer, the scaffold loop variant formed a tight band corresponding to well-folded monomer and the blunt end variant formed a ladder of bands, indicative of monomers and low order oligomeric states. This confirms that aggregates at higher MgCl_2_ concentrations consist of well-formed monomeric subunits rather than misfolded structures.

### Dependence of length of disordered strands on passivation

We next characterized the effect of the length of disordered regions on the efficacy at preventing aggregation. We synthesized a long scaffold loop variant by omitting all of the terminal staples on both the left and right sides of the blunt-ended variant, leaving ~30 nt scaffold loops **(Figure 3A, Supplementary figure 12)**. This longer scaffold loop variant yielded a higher proportion of well-folded monomers compared with the short scaffold loop variant across all MgCl_2_ concentrations tested (apart from the 5 mM sample mentioned previously), suggesting that longer scaffold loops were more effective than shorter scaffold loops at preventing aggregation **(Figure 3B, Supplementary figure 13)**. The highest yields were observed at 10 mM MgCl_2_ (88.2 ± 1.8%), which was similar to the highest yields obtained from the T_5_ brush variant. Like the short scaffold loop variant, there was a decline in the yield of monomeric subunits with increasing magnesium.

**Figure 3.**
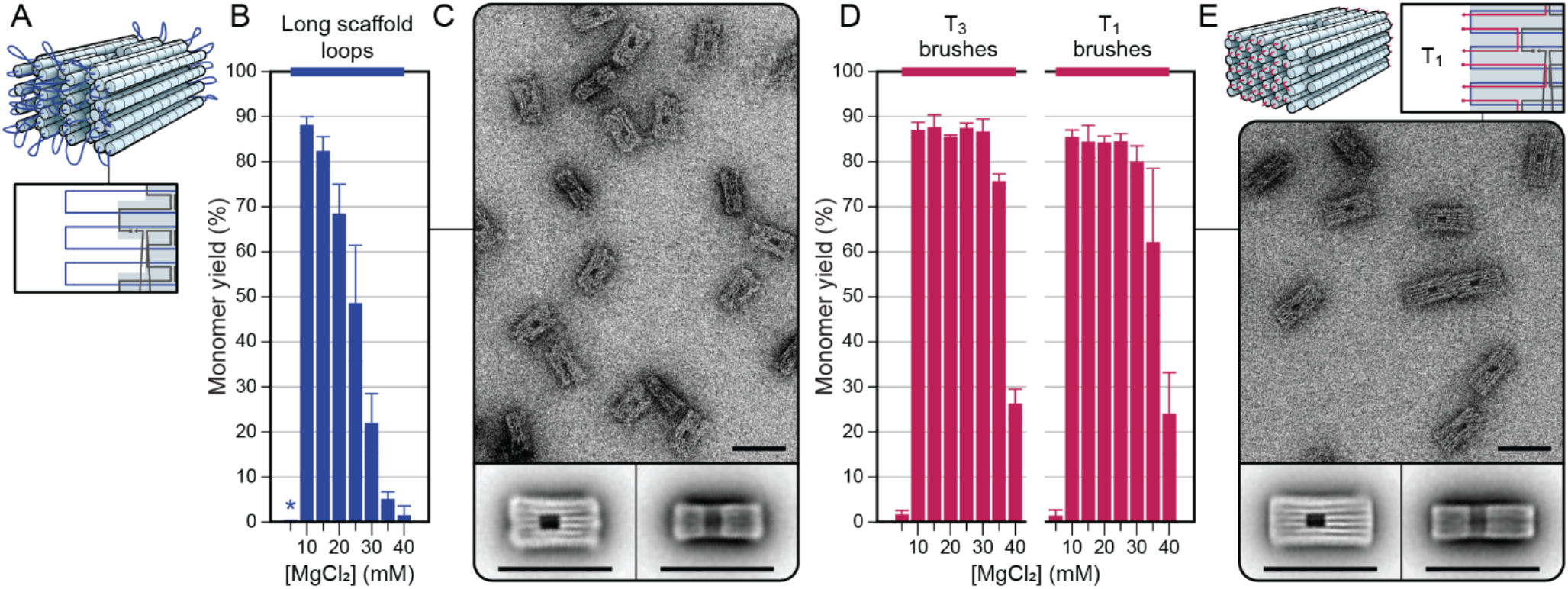
Effect of ssDNA length on aggregation prevention. A) Schematic of a design variant with longer (29 nt) scaffold loops. B) Monomer yields for the long scaffold loop variant at different concentrations of MgCl_2_. C) Exemplary TEM micrograph and 2D particle averages of the long scaffold loop variant. D) Monomer yields for the T_3_ and T_1_ brushed variants at different MgCl_2_ concentrations. E) Exemplary TEM micrograph and 2D particle averages of the T_1_ brush variant.

The long scaffold loop variants annealed in 20 mM MgCl_2_ appeared mostly well-formed, with approximately 5-10% of the structures appearing defective and very little aggregation evident in TEM micrographs **(Figure 3C, Supplementary figure 14)**. The length of this structure along the DNA helices appeared shorter than other variants, which was consistent with the design having fewer staples at the ends, and hence shorter double-stranded helices. The small number of aggregates that were visible in the micrographs appeared disordered, similar to the short scaffold loop variant, rather than linear, like the blunt-end variant. The fact that longer scaffold loops were more effective at passivation than shorter loops suggests that the entropic cost of inter-subunit aggregation with longer scaffold loops dominates over the enthalpic gain from forming additional base pairing interactions. It is also possible that the additional conformational flexibility in longer scaffold loops facilitate more intra-subunit base pairs than in shorter loops, further reducing the enthalpic gains from inter-subunit interactions and decreasing their chances of aggregating.

Since T_5_ brushes effectively prevented aggregation in up to 35 mM MgCl_2_, we reasoned that it would be informative to characterize the ability of shorter poly-T brushes to prevent base stacking. Thus, we synthesized additional T_3_ and T_1_ variants **(Supplementary figure 3)**. As with shorter scaffold loops, shorter poly-T brushes were less effective at preventing aggregation, however, T_3_ brushes performed almost as well as T_5_ brushes, with a decrease in monomer yield only observed in high MgCl_2_ (>30 mM) **(Figure 3D, Supplementary figure 15)**. Surprisingly, even structures with just a single thymine staple extension (T_1_) had monomer yields above 80% when annealed in buffer with MgCl_2_ between 10-25 mM **(Supplementary figure 16)**.

When PEG purified and resuspended in low magnesium buffer (5 mM MgCl_2_), both of the T_3_ and T_1_ brush variants appeared well-fomed in electron micrographs, with only a small amount of end-end stacking aggregation visible in the T_1_ variant **(Figure 3E, supplementary figures 17 & 18)**. However, when resuspended in high magnesium buffer (20 mM MgCl_2_), the T_1_ variant exhibited extensive base-stacking aggregation that was not evident in the same sample when examined via AGE **(Supplementary figure 19A**,**B)**. The reason for the discrepancy between AGE and TEM results for this variant is unclear, but is likely due to the differences in sample treatment and preparation. Preparation for TEM grid staining is slower, allowing time for aggregates to form, while AGE may break up weakly-bound aggregates while migrating through the gel matrix. We also noticed that the temperature of the electrophoresis buffer affected the strength of secondary bands, with colder buffer displaying more evidence of aggregation in high Mg samples **(Supplementary figure 19C)**.

### Passivation via staple extensions with sequences other than poly-T

The length-dependence of scaffold loops suggests that disordered regions create an entropic barrier to aggregation that dominates over potential binding mechanisms including random base pairing interactions. Given this, the necessity for the near exclusive use of the poly-T as extensions to prevent unwanted base-stacking, may be unwarranted and unecessarily limit our versatility in controlling interactions between DNA origami structures. Indeed, LaBean and colleagues mentioned that in their hands A_4_ loops were as effective as T_4_ loops in preventing aggregation.^[24]^

AGE and densitometry measurements indicated that poly-A extensions were simlarly effective at preventing DNA origami aggregation as the same length poly-T extensions at 10-15 mM MgCl_2_, exhibiting monomer yields as high as 90%, but were less resiliant to preventing aggregation at higher MgCl_2_ concentrations **(Figure 4A, Supplementary figures 20-22)**. Ordered linear polymers ranging in length from 2-20 subunits were visible in TEM micrographs when the A_1_ brushed variants were annealed in 20 mM MgCl_2_ and resuspended in 5 mM MgCl_2_ **(Figure 4B, Supplementary figure 23)**. This polymerisation occurred to a much lesser extent in A_3_ brush variants, and was completely absent in the A_5_ variant **(Supplementary figures 24**,**25)**. Like the T_1_ variant, the A_1_ variant exhibited increased aggregation when resuspended post-purification in buffer with high Mg concentration, and also when electrophoresed at lower temperatures **(Supplementary figure 26)**.

**Figure 4.**
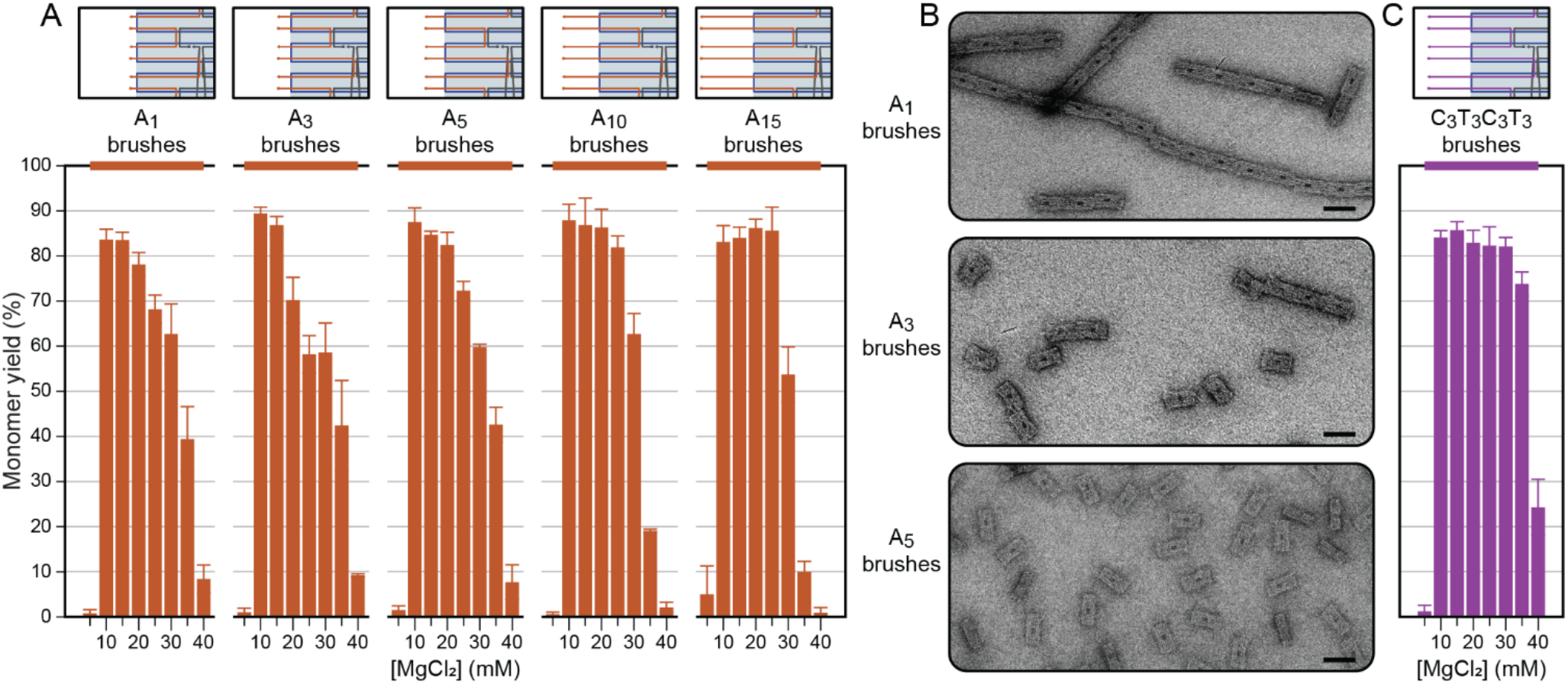
DNA origami structures passivated with brushes of various lengths and nucleotide composition. **A)** Monomer yields of adenine brushes of length 1, 3, 5, 10 and 15 nt annealed in different MgCl_2_ concentrations. **B)** Electron micrographs of A_1_, A_3_ and A_5_ variants annealed in 20 mM MgCl_2_ and buffer exchanged into 5 mM MgCl_2_. Scale bars are 50 nm. **C)** Monomer yields of cytosine/thymine (C_3_T_3_C_3_T_3_) variant.

DNA origami structures with longer poly-A (A_10_ or A_15_) extensions performed similar to T_5_ extensions up to 30 mM MgCl_2_ where monomer yields were higher with T_5_ extensions (Figure 4A, **Supplementary figures 27-30**). Monomer yields were similar between A_10_ and A_15_ extensions across all MgCl_2_ concentrations, suggesting maximal passivation was achieved with a 10 nt extension.

Testing the suitability of the other two nucleotides, cytosine and guanine, for preventing aggregation is complicated by the propensity of poly-C and poly-G strands to form secondary structures such as the i-motif and G-quadruplex, respectively,^[25, 26]^ Thus, to investigate the use of poly-C strands, we designed a C_3_T_3_C_3_T_3_ variant in which staple extensions on the DNA origami structure consisted of CCCTTTCCCTTT on one side or TTTCCCTTTCCC on the other with the thymines proximal to the DNA origami (Supplementary figure 3). These sequences were chosen because they do not form secondary structures at the temperature and pH that all our experiments were conducted (25°C & pH 8.0).^[27]^ This variant was resistant to aggregation with near maximal yields of monomers (>80%) between 10-30 mM as quantified with AGE and densitometry **(Figure 4C, Supplementary figure 31)**.

### A novel strategy for preventing aggregation: Orthogonal end-capping

To date, disordered single-stranded DNA is the only structural mechanism shown to be effective at passivating arrays of exposed blunt-ends in DNA origami structures. Consequently, the border between DNA origami structures and solution is often ill-defined, akin to a polymer brush.^[28]^ Moreover, such disordered regions act like a entropic springs, imparting a repelling force that limits how close adjacent structures can be located (ref PolyBrick). We therefore developed a novel strategy for preventing inter-subunit base-stacking that does not require disordered single-stranded DNA.

This strategy involved shielding the blunt ends of DNA origami structures with a single layer of double-stranded DNA. The strategy was achieved by routing the scaffold strand in such a way that long scaffold loops connect helices across the terminus of the origami structure, perpendicular to the direction of the helices in the bundle. These long scaffold loops were hybridized to complementary staple strands, creating double-stranded helices orthogonal to the helices in the core structure and capping the ends to prevent intermolecular base stacking. We therefore refer to this strategy as “orthogonal end-capping” (OEC).

We implemented the orthogonal end-capping strategy on a 42-helix bundle with the same layout as the previous DNA origami designs to compare its effectiveness in preventing aggregation (**Supplementary figure 32)**. There were 6 orthogonal scaffold loops, each spanning the 6 columns of 8 helices at the end of the honeycomb lattice, forming a horizontal zig zag pattern **(Figure 5A)**. Each scaffold loop was 46 nt, of which 42 nt were hybridized to an orthogonal staple, leaving a 2 nt unpaired spacer at each end to allow some flexibility and prevent possible base stacking along the edges of the orthogonal helices. The length of orthogonal helices was designed to correspond to the span of the 8 helices in each of the 6 columns at the end of the honeycomb lattice design **(Supplementary figure 33)**.

**Figure 5.**
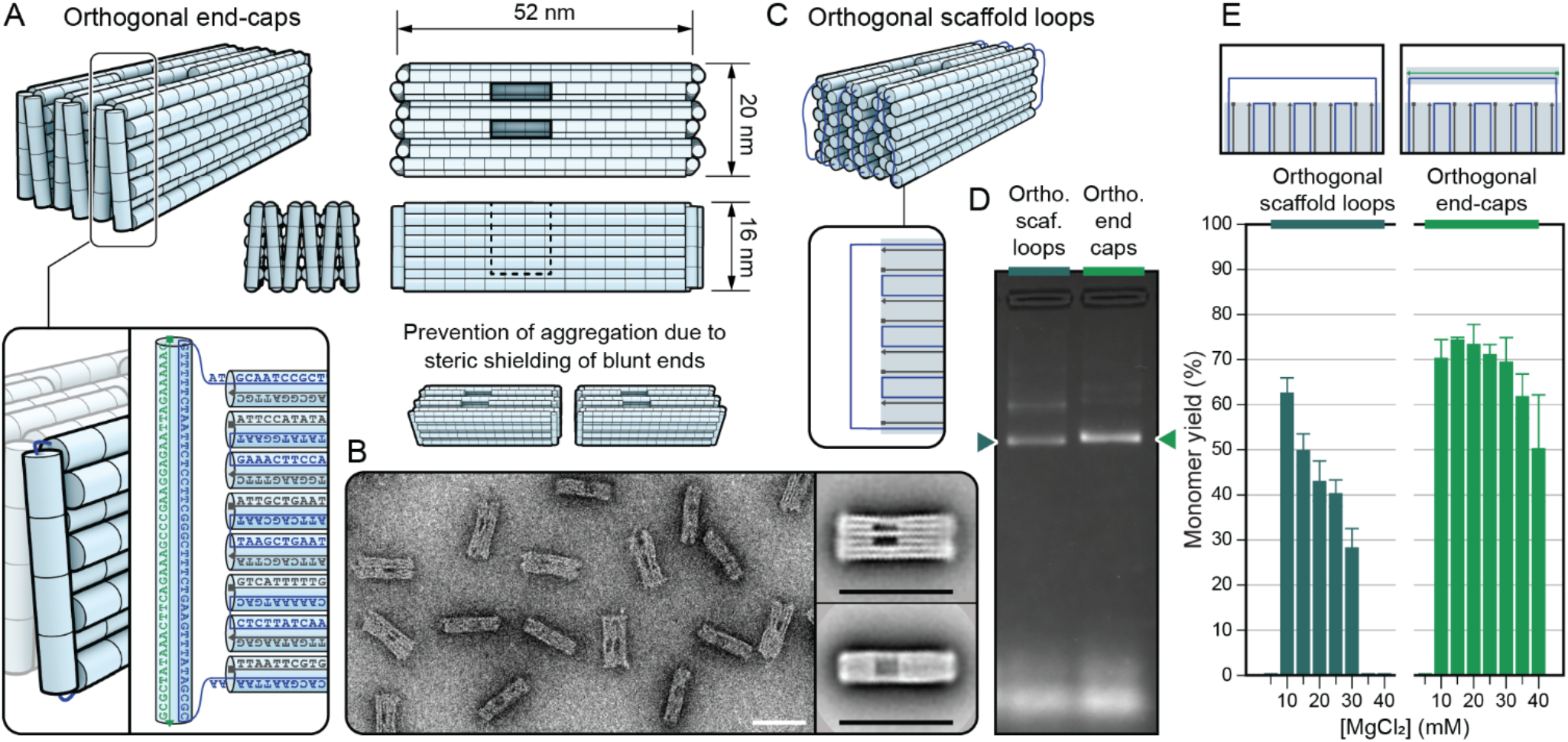
Orthogonal end-capping to prevent aggregation. **A)** Design schematics of the orthogonal end-capped DNA origami structure. **B)** An exemplary TEM micrograph and 2D particle averages for the orthogonal end-capped design. Scale bars are 50 nm. **C)** Design of the orthogonal scaffold loop variant, in which the orthogonal staples were omitted. **D)** Agarose gel electrophoresis results of the two variants annealed at 20 mM MgCl_2_, with arrows indicating well-folded monomer bands. **E)** Monomer yields of the two variants annealed with different concentrations of MgCl_2_.

We synthesized the OEC structure using the same protocols as used for the previous designs and analysed them using TEM and AGE. Under TEM, the structures appear mostly well-folded and monomeric, with ~10% of the particles exhibiting obvious defects, and only ~3% of the particles appearing in dimers with no larger aggregates **(Figure 5B, Supplementary figure 34)**. When we omitted the end-cap staples creating long orthogonal scaffold loops **(Figure 5C)**, a greater proportion of structures appeared aggregated, indicating benefit in capping structures with orthogonal helices over disordered scaffold loops for preventing aggregation **(Supplementary figure 35)**.

We quantified the yield of monomers in both variants with AGE (**Figure 5D,E**, **Supplementary figures 36**,**37**). The OEC structure had a maximal yield of monomers at MgCl_2_ concentrations between 10 – 30 mM MgCl_2_, thus exhibiting a similar resilience to MgCl_2_ as the T_5_ brush design, but with a lower maximal yield of monomers of around 70%. The OEC variant also exhibited several faint bands that migrated at a slower rate than the bright main band. Some of these bands disappeared when purified and resuspended into buffer with low MgCl_2_, with the decrease in brightness from the faint bands commensurate with an increase in the brightness of the main band **(Supplementary figure 38)**. Given that previous experiments showed well-folded but aggregated origami structures could be separated into monomers in low-magnesium buffer, we concluded that this slower band consisted of misfolded structures, rather than aggregates, a result also observed in TEM micrographs, thus explaining the lower maximal yield of monomers as measured by densitometry.

The orthogonal scaffold loop variant had a lower monomer yield compared to the OEC variant across all MgCl_2_ concentrations. Its aggregation showed a similar trend to that of the long scaffold loop variant examined earlier, with its monomer yield highly dependent on MgCl_2_ concentration and exhibiting a much higher proportion of aggregated monomers that could be separated in low magnesium buffer. This indicates that the effective prevention of aggregation seen in the OEC can be attributed to the double-stranded end caps, which minimise base-stacking as well as base-pairing at the ends of the structures.

### Controlling multi-subunit DNA origami assembly

Although less effective at preventing aggregation than poly-N (e.g. poly-T or poly-A) staple extensions, the distinct sequences in scaffold loops **(Figure 6A(i))** facilitate controlled assembly to synthesize multi-origami assemblies.^[29]^ This multimer assembly can be achieved in two main ways. First, by adding single-stranded linkers that are complementary to the scaffold loops of two structures and hence can form a double-stranded bridge between them once annealed **(Figure 6A(ii))**. The advantage of this method is that assembly of the structures can be triggered at a specific time in the experiment by the addition of the linkers. The disadvantage is that the structures must be purified from excess staples before assembly.

**Figure 6.**
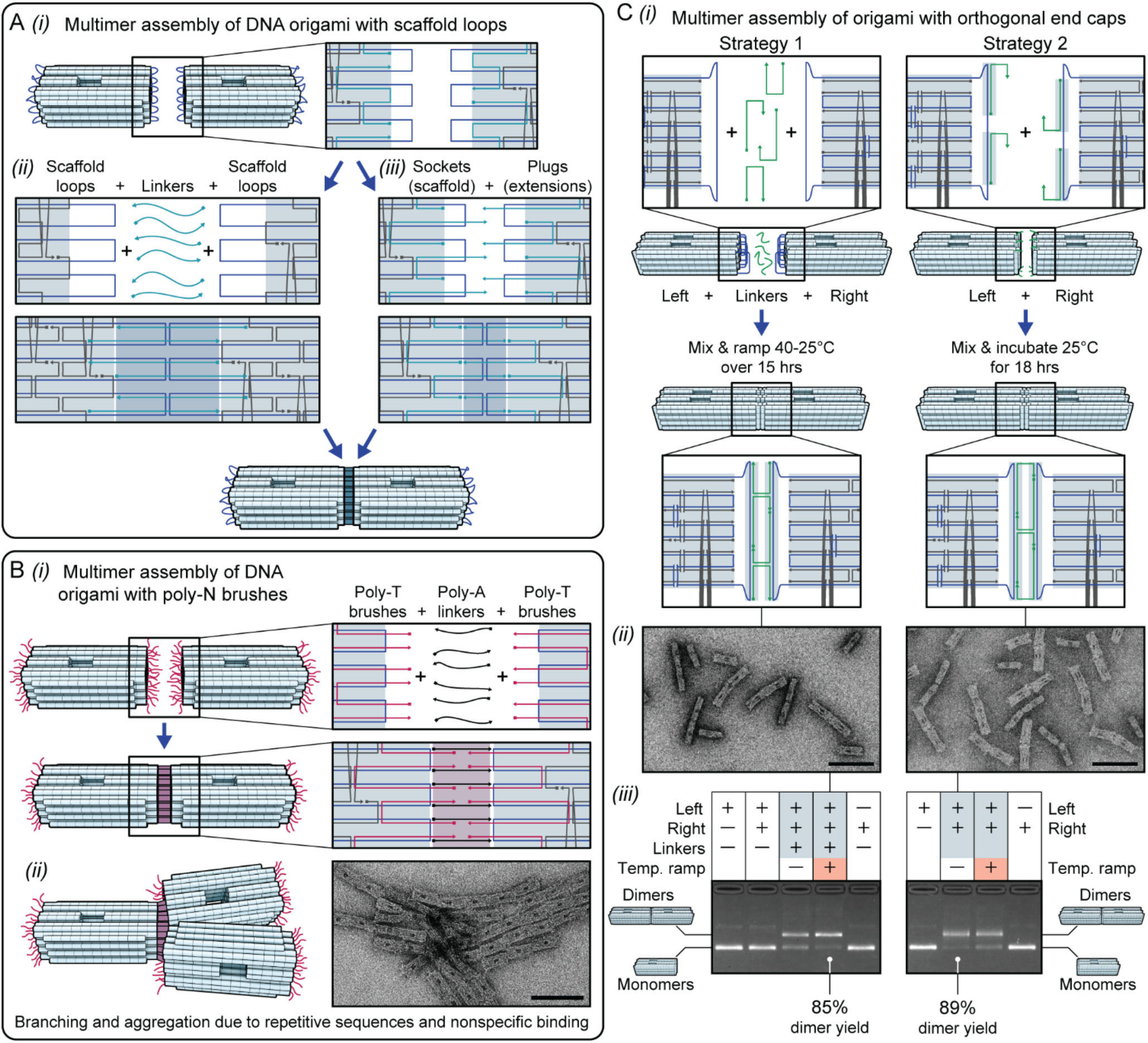
Multimeric assembly of DNA origami structures with different passivation strategies. **A)***(i)* Structures with scaffold loops can be precisely assembled via the post-purification addition of *(ii)* linkers that traverse the scaffold loops of multiple subunits, or *(iii)* via the formation of alternating short staple extensions (plugs) and staple truncations that reveal scaffold (sockets). **B)***(i)* Designs that feature poly-T staple extensions can be assembled with similar structures by adding poly-A linker strands, but this can lead to *(ii)* misaligned, branched and aggregated assemblies. **C)***(i)* Schematics of two assembly strategies for the orthogonal end-capped structures. *(ii)* Exemplary TEM micrographs of each strategy. *(iii)* AGE results for each strategy, with yields of dimers calculated from gel densitometry. Scale bars in TEM micrographs are 100 nm.

The second method of assembly is similar, but the linker strands are annealed with the origami structures as they are folded, leaving short single-stranded extensions (plugs) and complementary regions of unpaired scaffold (sockets) **(Figure 6A(iii))**. Structures are therefore directed to bind to each other simply by mixing them together without the need for an additional step adding the linkers. Both scaffold loop binding methods have been shown to be effective in creating high yields of multimeric assemblies of DNA origami subunits.^[3, 28, 30-32]^

Poly-T (or indeed any repeating sequence) extensions provide less control for multimeric assembly because they present multiple identical binding sequences at the binding interface **(Figure 6B(i))**. These extensions can be bound to complementary linkers to induce assembly with other origami subunits, but each linker can bind at multiple places on the binding interface, and this can lead to branched filament assembly or large disordered aggregates. We attempted multi-subunit assembly of DNA origami with poly-N brushes by adding poly-A linkers to poly-T-decorated structures and *vice versa*. For all variants tested we observed large, branched aggregates and fibrils in TEM micrographs and a failure of any structures to migrate during electrophoresis **(Figure 6B(ii), Supplementary figures 39-41)**. End-to-end polymeric assembly could be somewhat assisted by thermal ramping of the structures after adding the linkers, but misalignment of adjoining monomers remained a problem.

We have demonstrated that OECs can be as effective as poly-N staple extensions for preventing aggregation (Figure 4B,E). Yet OECs also present a unique sequence that can be harnessed to control multi-subunit assembly without compromising control of unwanted aggregation, thus providing benefits of scaffold loops and poly-N brushes. We devised two strategies to achieve controlled multimeric assembly of OEC origami structures **(Figure 6C(i))**. In each strategy, ‘left’ and ‘right’ variants were designed, each with one ‘passive’ side with OECs as previously described, and one ‘active’ side modified to facilitate multi-subunit assembly. These variants were synthesized and purified separately, incubated together in equal ratio, and examined via TEM and AGE to determine yields of dimer.

Strategy 1 involved folding the left and right variants with active sides comprising orthogonal scaffold loops with no staples. After purification from excess staples, left and right variants were mixed together in the presence of a excess of linker strands designed to bridge the orthogonal helices between them. Strategy 2 involved folding the left and right variants with 5 nt plugs and sockets. These did not need linker strands, with assembly initiated simply by mixing.

TEM micrographs for both strategies showed high yields of well-formed and correctly aligned dimers, and no disordered aggregates **(Figure 6C(ii), Supplementary figure 42)**. Analysis by AGE demonstrated that in each strategy, neither subunit formed dimers independently of its complement, and in the case of strategy 1, a mixture of left and right monomers required linkers to form dimers **(Figure 6C(iii))**. For each strategy we conducted two incubation conditions; one was incubated at room temperature overnight, and one underwent thermal ramping from 40 °C to 25 °C over 15 hours. Dimer yields were calculated from gel densitometry. For strategy 1, temperature ramping created the highest dimer yield of 85%, while for strategy 2, overnight incubation at room temperature was more effective, with a dimer yield of 89%. These high yields of correctly assembled dimers illustrate the unique suitability of the OEC passivation strategy for both prevention of aggregation in diverse conditions and for precise multimer assembly.

## Conclusion

In this study, we present the first systematic exploration of various design techniques for the passivation of DNA origami structures. This was achieved by designing a core structure that presented a consistent number of coplanar DNA ends (48-helix bundle), thereby providing a large surface for base-stacking while also typifying DNA origami designs. The core structure was then synthesized with several variations, each with a different method of passivation. These included a blunt-ended control variant, several with methods for passivation already widely used in DNA origami design (scaffold loops, poly-T brushes) and some novel or less-frequently used methods (poly-A brushes, orthogonal end-caps). All variants were assessed for their efficacy at preventing aggregation in a range of configurations and MgCl_2_ concentrations, and their ability to mediate controlled multi-subunit assembly.

We found that poly-T brushes produced the highest yield of well-formed monomeric subunits but provide minimal control for mediating multi-subunit assembly. Moreover, poly-A or poly-C brushes could be similarly effective as poly-T brushes at preventing aggregation in certain chemical conditions, despite the ubiquity of thymines for passivation of DNA origami structures. Scaffold loops appeared to be effective at preventing aggregation from base-stacking but could also cause aggregation by introducing inter-subunit base-pairing and thus were less effective than poly-T brushes for passivation. However, unlike poly-T brushes, scaffold loops enable control of multi-subunit assembly. We also introduced a novel OEC passivation method, which we demonstrate has similarly efficacy as poly-N brushes for passivation but simultaneously enables controlled multi subunit assembly thus providing the benefit of both poly-N brushes and scaffold loops. We also demonstrated that longer single-stranded extensions are better than shorter extensions for preventing aggregation, presumably due to a larger repulsive entropic force between subunits. This applied to both scaffold loops as well as poly-N staple extensions.

In this study, we used a single core DNA origami structure to characterise different passivation methods. To generalise these findings further, it will be necessary to test passivation strategies on different structures since it is possible that their relative effectiveness at preventing aggregation may depend on the size, cross-sectional shape, or lattice arrangement of the origami structure. It is also unclear on whether the OEC strategy can be practically implemented on any DNA origami structure. We attempted to synthesize two variants in which the orthogonal DNA helices ran horizontally across the end of the structure instead of vertically **(Supplementary figures 43-46)**, but both variants suffered from poor folding yields in all synthesis conditions tested. Thus, while further exploration is necessary to determine the adaptability of the OEC strategy to other designs, this study represents a significant step towards the development of effective and versatile methods for the prevention of aggregation in DNA origami structures.

## Materials and Methods

### DNA origami design, synthesis and purification

DNA origami structures were designed using Cadnano^[33]^ and Cando^[34]^ iteratively. Scaffold strands (10 nM, M13mp18, Bayou Biolabs) were mixed with staple strands (100 nM, Integrated DNA Technologies) in 1× TE buffer (10 mM Tris, 1 mM EDTA) with varying concentrations of MgCl_2_ (20 mM unless otherwise specified). Mixtures were annealed in a thermocycler (Bio-Rad) by first heating to 65 °C for 15 min, then slowly cooling from 60 °C to 44 °C at 6 min/0.1 °C, then holding at 25 °C. Agarose gel electrophoresis was done in 2% agarose gels with 0.5× Tris-Borate-EDTA buffer with 6 mM MgCl_2_ and RedSafe™ Nucleic Acid Staining Solution (Intron), and electrophoresed at 70 V for 3 hours at room temperature. Gels were imaged with a UV transilluminator GelDoc (BioRad) and gel densitometry was done using GelAnalyser.^[35]^ Purification from excess staples was performed using PEG purification^[23]^, with resuspension into 1× TE buffer with varying concentrations of MgCl_2_ (5 mM unless otherwise specified).

### Imaging and analysis

PEG purified DNA origami samples were drop cast onto carbon/Formvar-coated copper grids (Ted Pella) which were first glow-discharged for 60 s and given 4 min to adsorb. After excess sample was wicked off, a drop of 2% aqueous uranyl formate was applied to the grid and wicked off immediately. Imaging was performed on an Tecnai G2 20 transmission electron microscope (FEI) in bright field mode at 200 kV. 2D particle averaging was done using Relion.^[36]^ Measurement was performed using ImageJ.^[37]^

### Multimeric assembly

T_5_, A_5_ or A_10_ brushed structure variants were assembled into polymers by mixing 5 nM origami with A_15_, T15 or T_21_ linker strands respectively in 10× stoichiometric excess for each binding site (48 per origami) and incubating either 18 hours at room temperature or by thermal ramping from 40 °C to 25 °C at 6 min/0.1 °C. Orthogonal end-capped structures were assembled into dimers by mixing left and right variants (5 nM each) and incubating as described in the main text. For scheme 1, linkers were added in 100× stoichiometric excess prior to incubation. All assembly reactions took place in 1× TE buffer with 10 mM MgCl_2_.

## Acknowledgements

The authors thank the facilities of Microscopy Australia at the Electron Microscope Unit, Mark Wainwright Analytical Centre, UNSW Sydney. This research was supported by the Australian Research Council Centre of Excellence in Synthetic Biology (Grant ID CE200100029)

## Notes

### Competing Interest Statement

The authors have declared no competing interest.

