## Supplementary Figure for "Passivating blunt-ended helices to control monodispersity and multi-subunit assembly of DNA origami structures"

### Design-based strategies for preventing aggregation of DNA origami

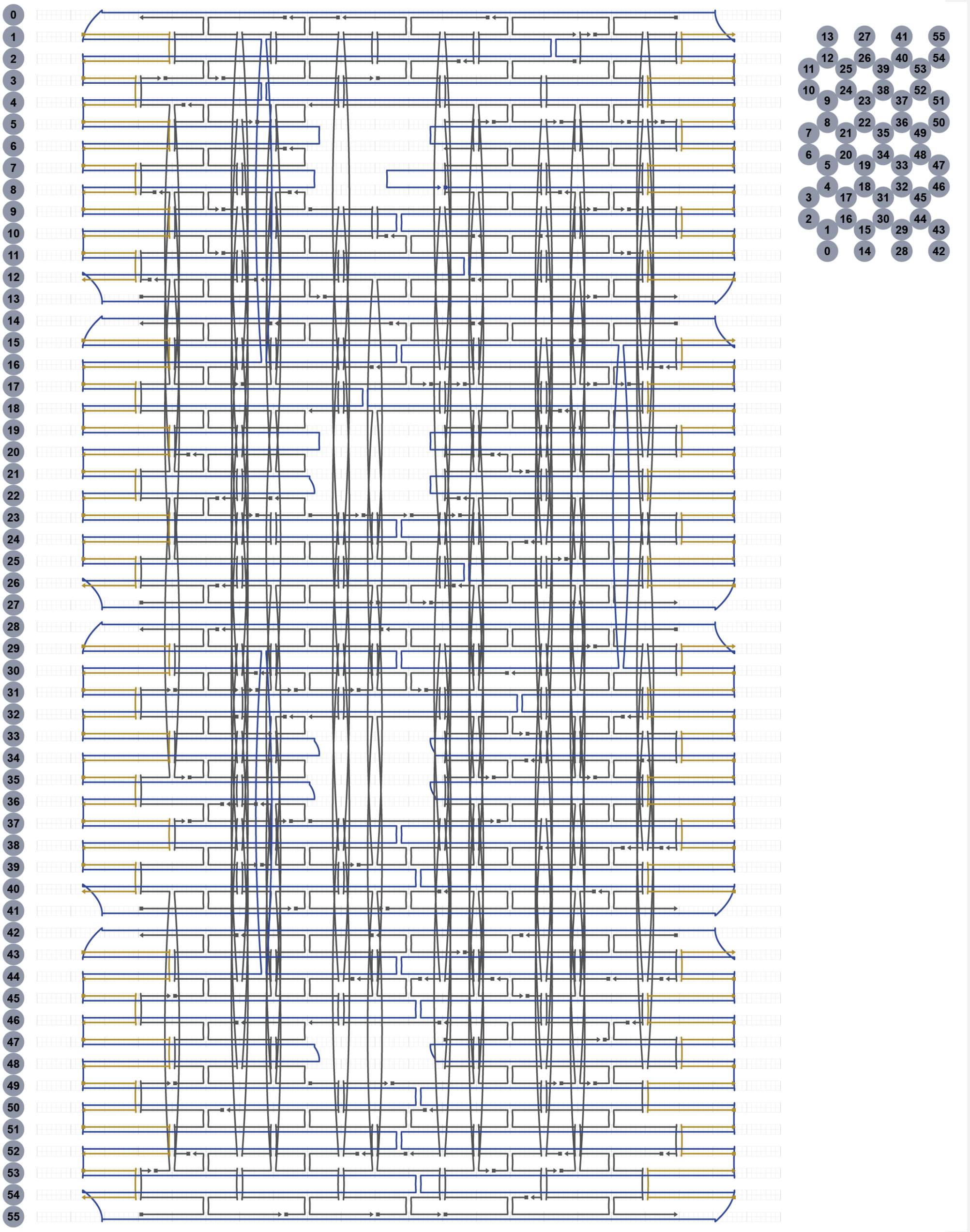


Supplementary figure 1 – Strand schematic layout of blunt-end variant

**Commented [LL1]:** Should we say Cadnano schematic?

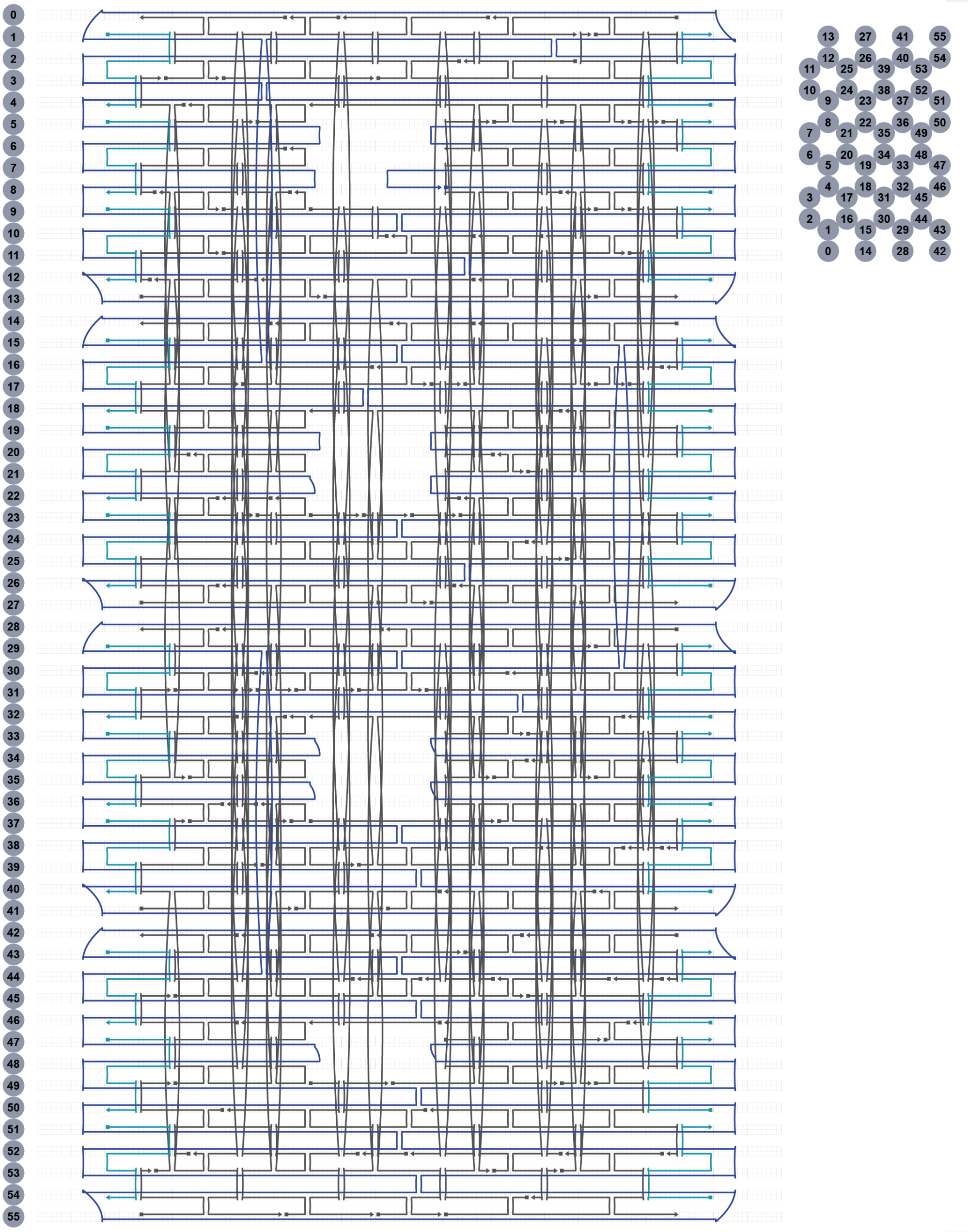


Supplementary figure 2 - Strand schematic layout of short scaffold loop variant

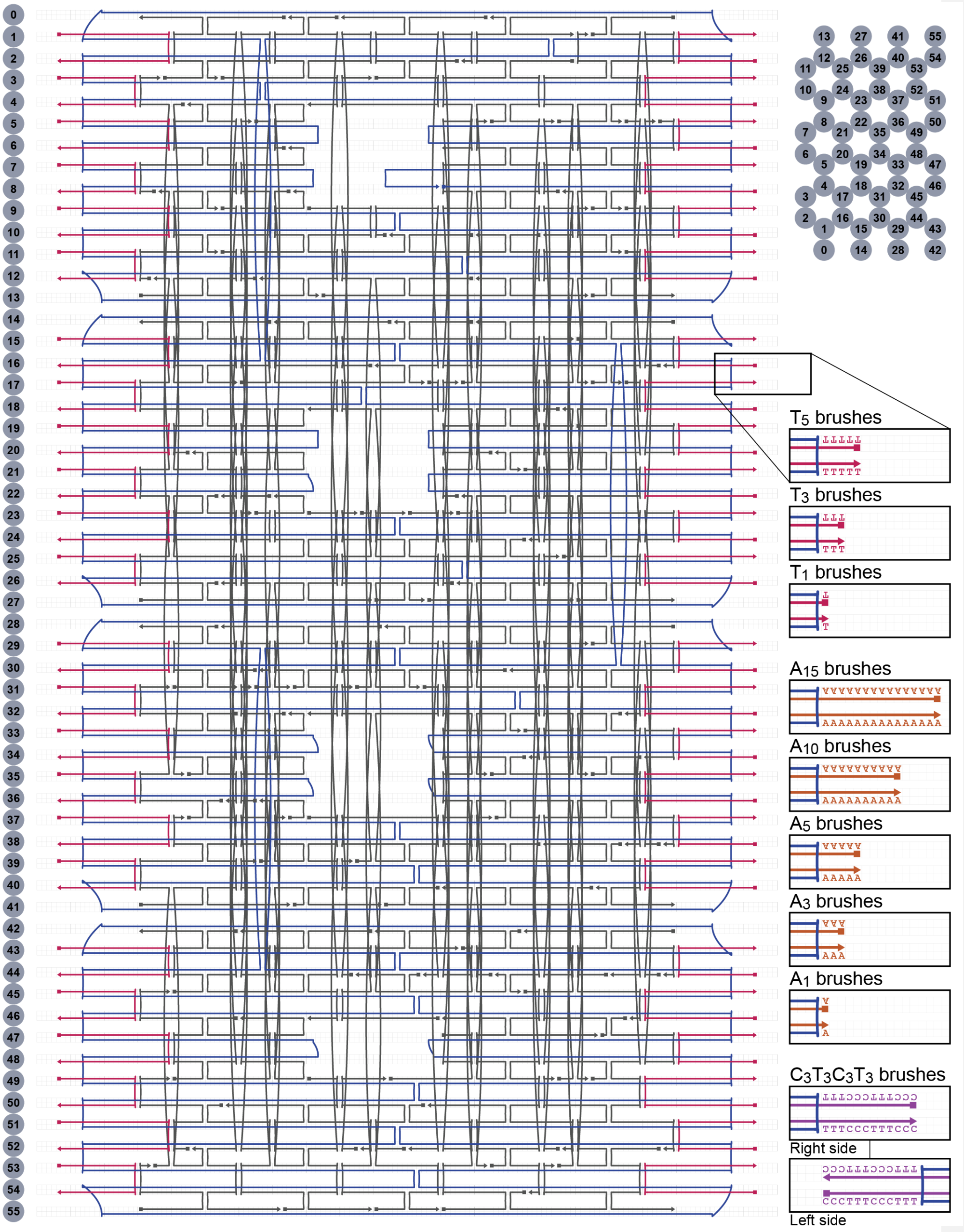


Supplementary figure 3 - Strand schematic layout of poly-N brush variant

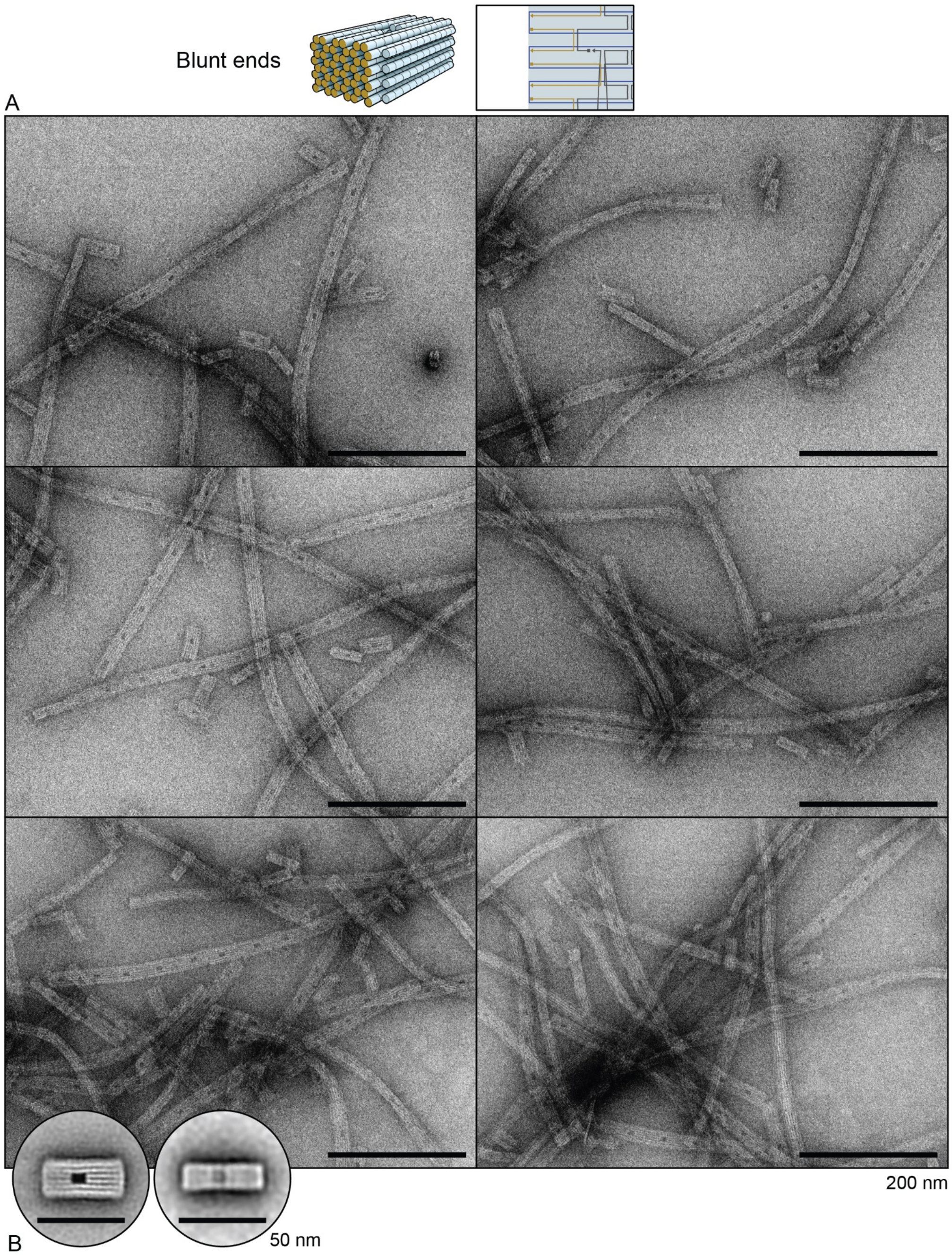


Supplementary figure 4 – Transmission electron micrographs of blunt-ended variant. **A)** Representative micrographs. **B)** Two-dimensional particle averages.

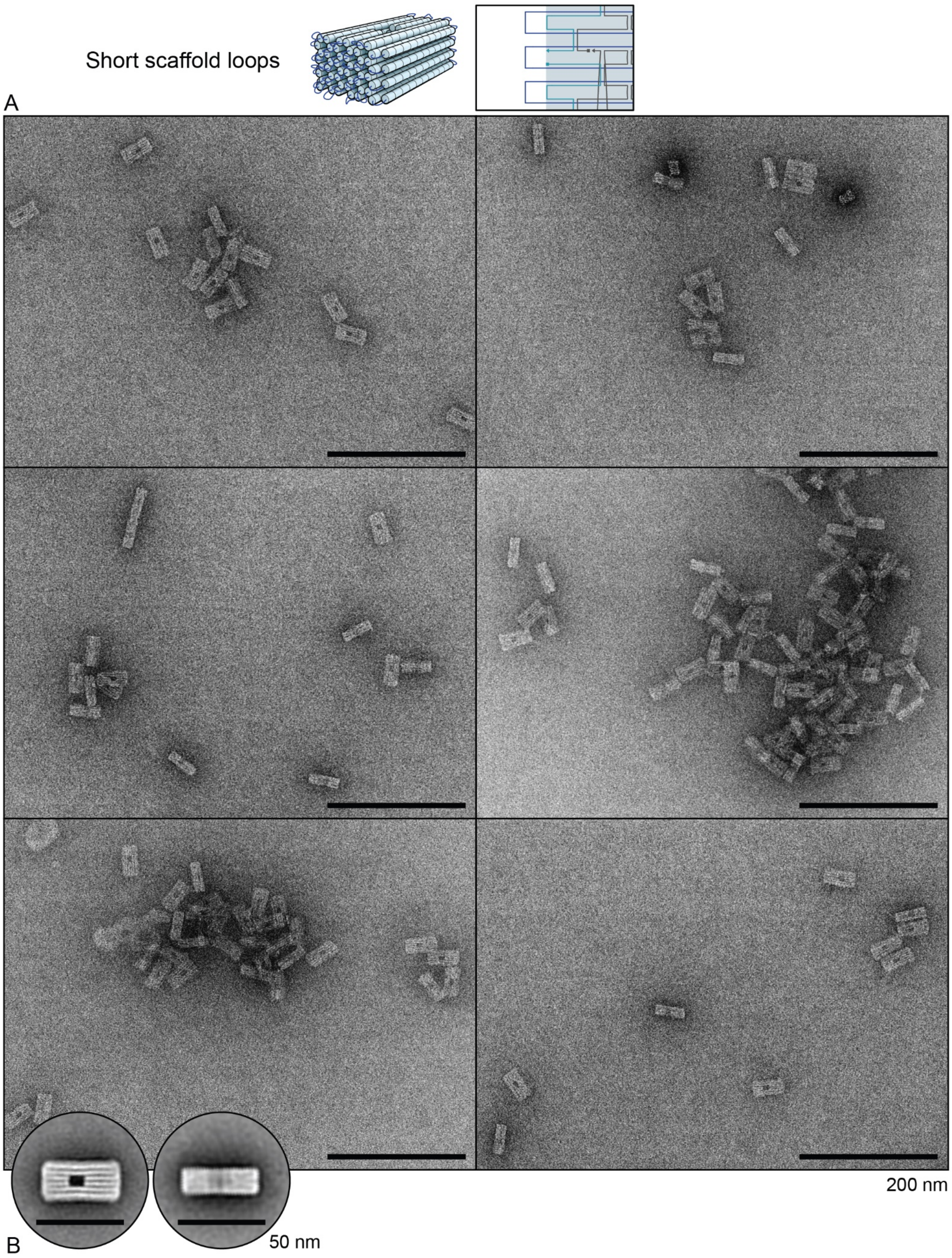


Supplementary figure 5 – Transmission electron micrographs of short scaffold loop variant. **A)** Representative micrographs. **B)** Two-dimensional particle averages.

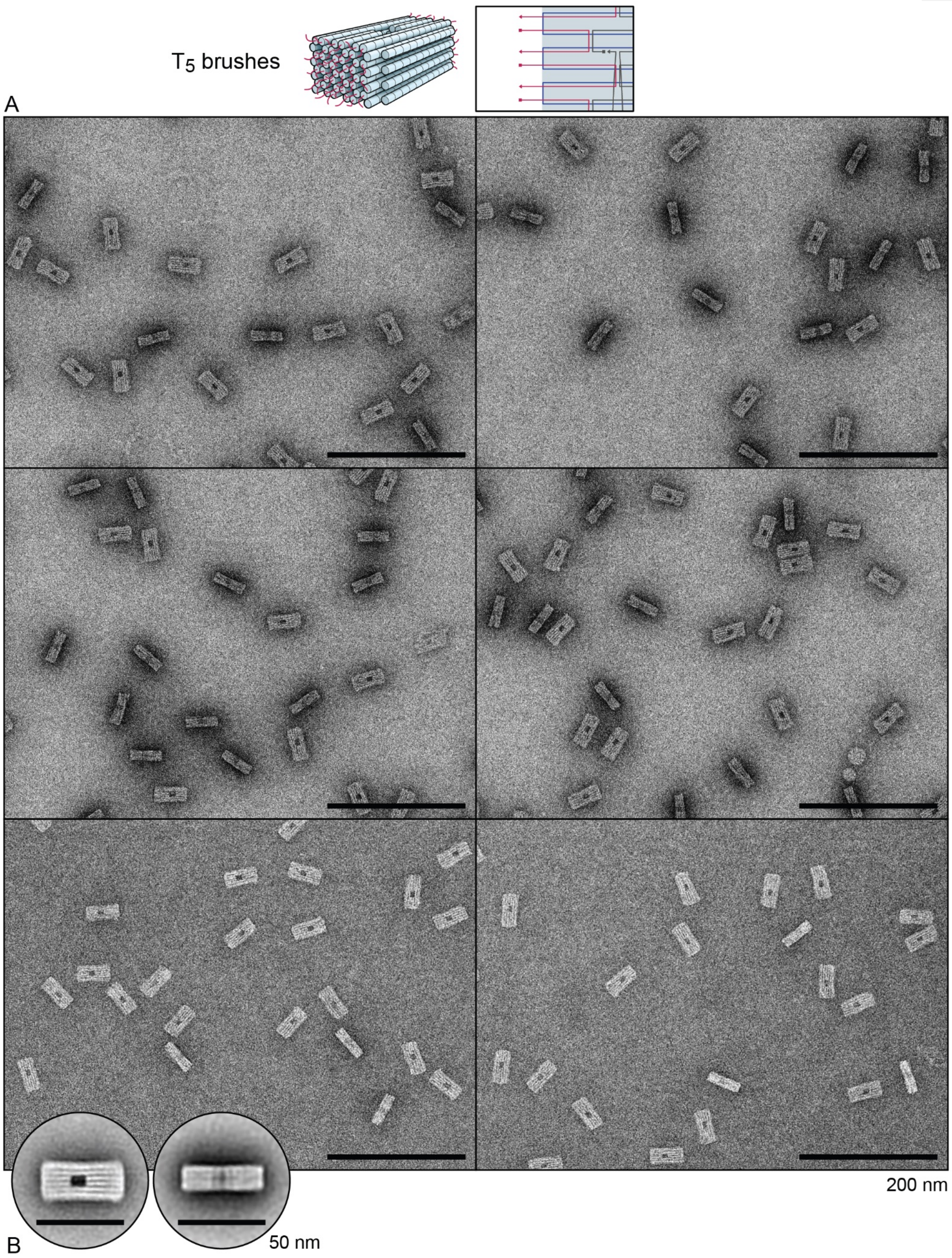


Supplementary figure 6 – Transmission electron micrographs of $T_5$ brush variant. **A)** Representative micrographs. **B)** Two-dimensional particle averages.

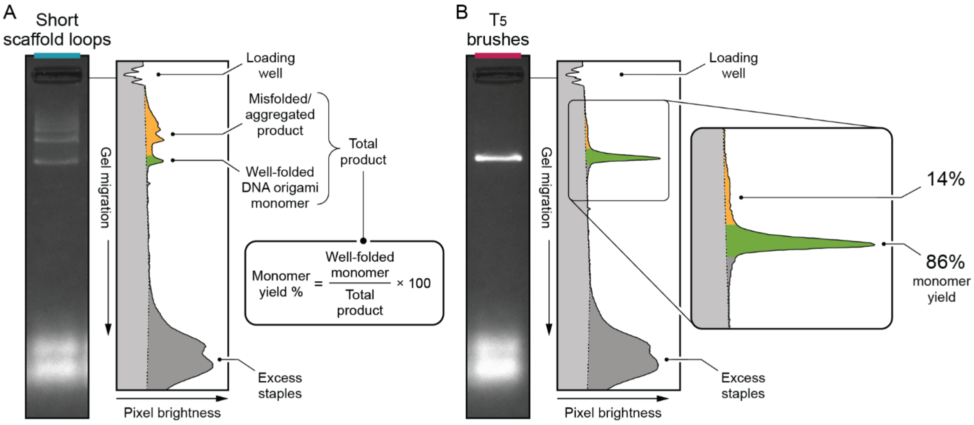


Supplementary figure 7 – Calculation of monomer yields from agarose gel densitometry. **A)** Short scaffold loop variant. **B)** $T_5$ brush variant.

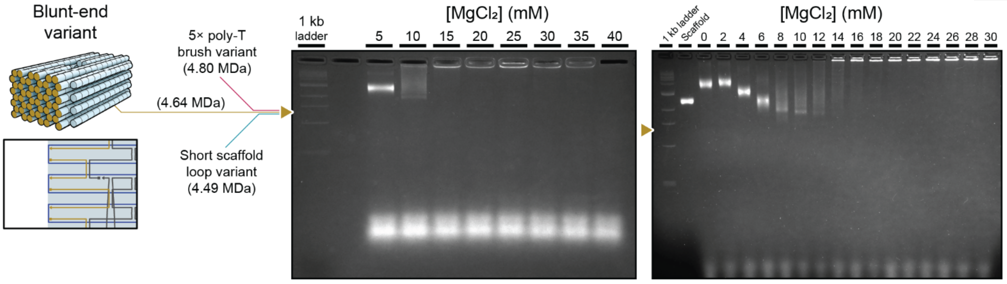


Supplementary figure 8 – Agarose gel electrophoresis results for the blunt-end variant. The triangle indicates the expected migration distance of well-formed monomeric structures, which was calculated by comparing the migration distances and molecular weights of other variants with well-formed monomers based on the migration rate of pacified DNA origami structures below.

Commented [LL2]: Correct?

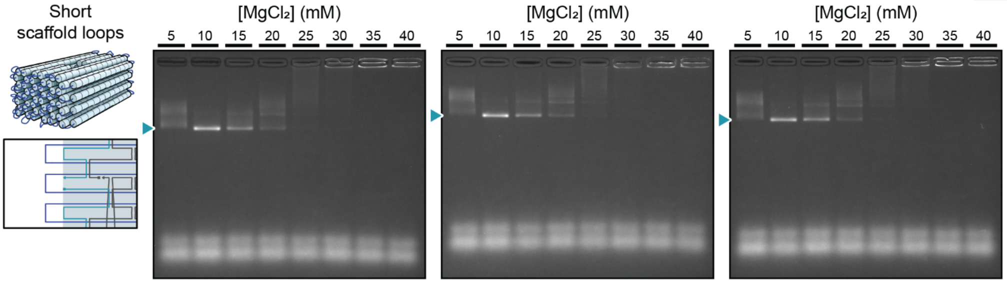


Supplementary figure 9 – Agarose gel electrophoresis results (3 replicates) for the short scaffold loop variant. The triangle indicates the migration distance of well-formed monomeric structures.

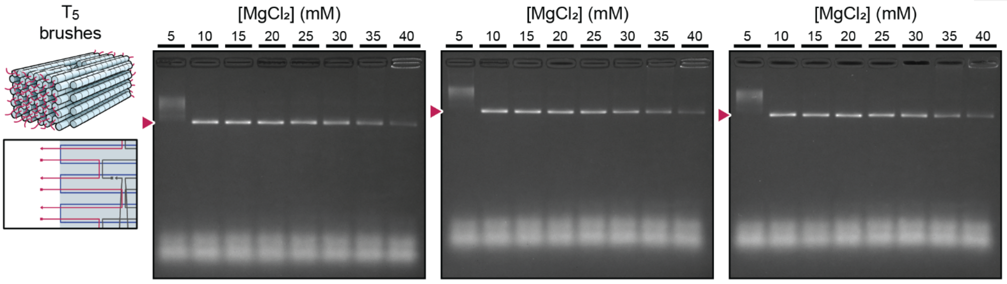


Supplementary figure 10 – Agarose gel electrophoresis results (3 replicates) for the $T_5$ brush variant. The triangle indicates the migration distance of well-formed monomeric structures.

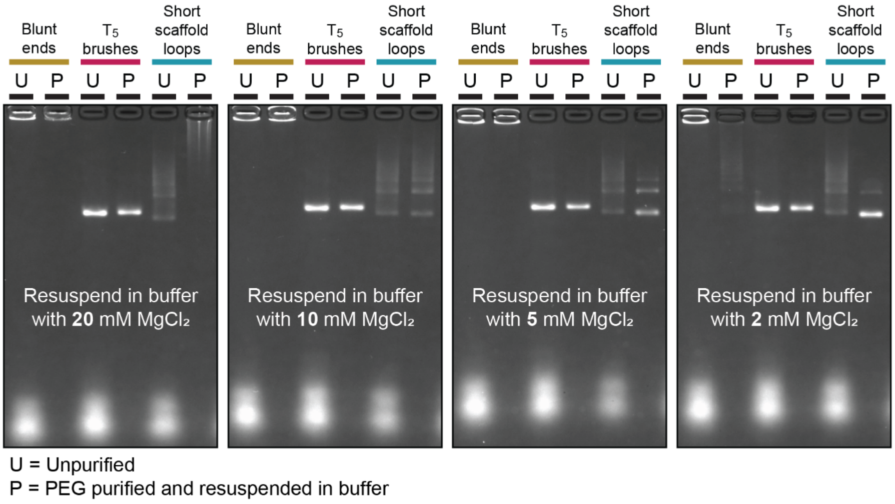


Supplementary figure 11 – Agarose gel results for DNA origami variants before and after purification and buffer exchange into lower $MgCl_2$

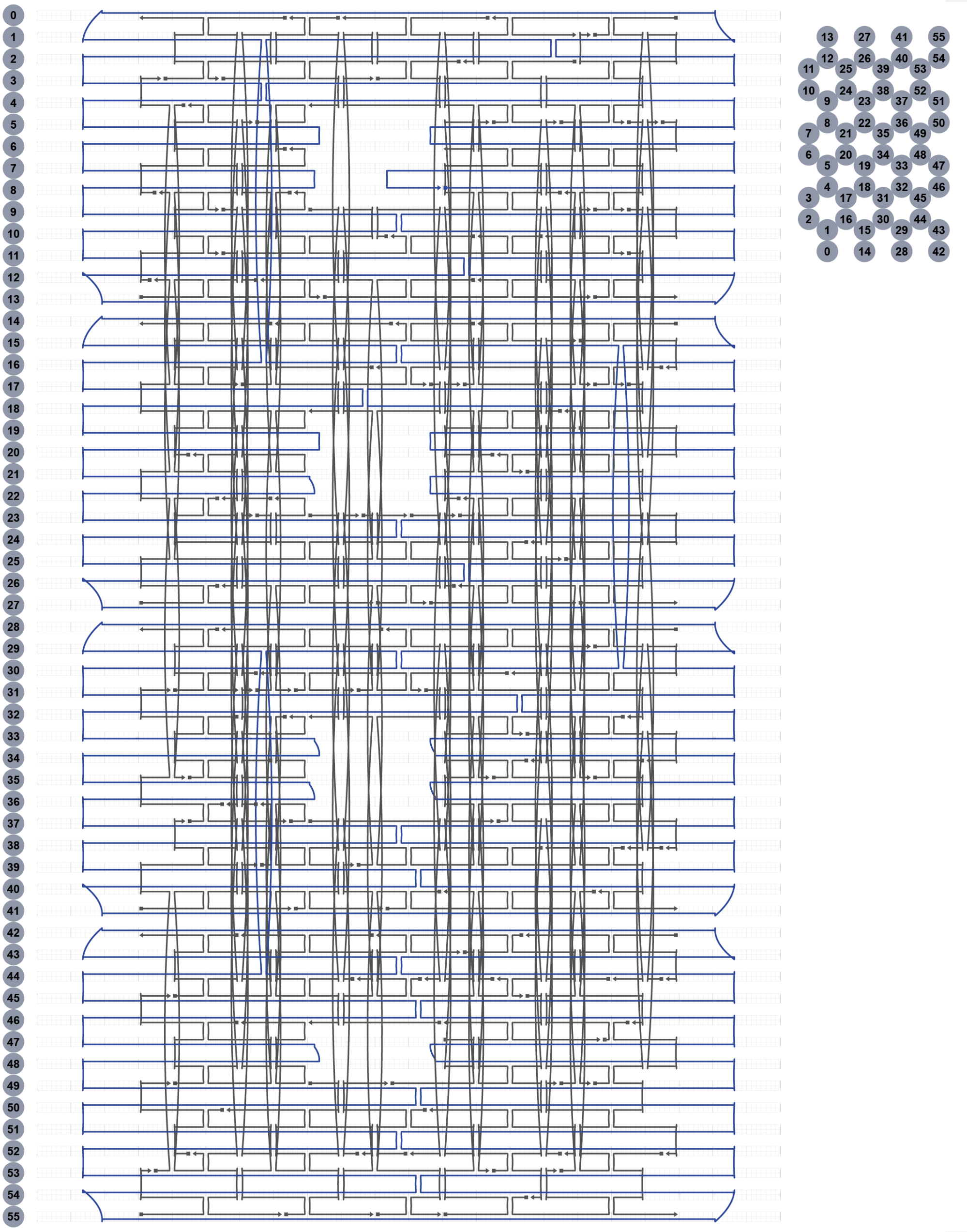


Supplementary figure 12 - Strand schematic layout of long scaffold loop variant.

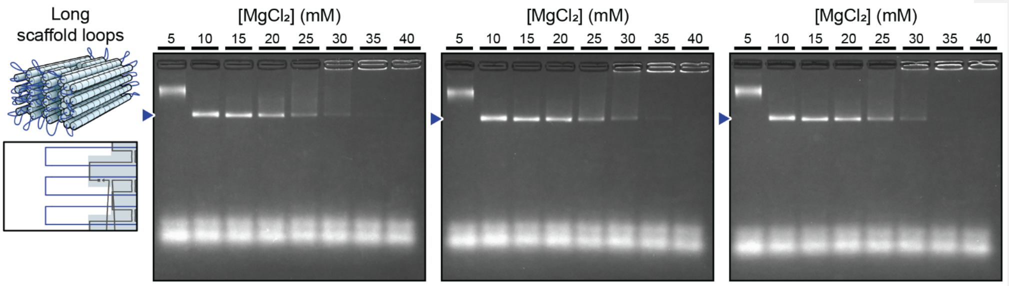


Supplementary figure 13 – Agarose gel electrophoresis results (3 replicates) for the long scaffold loop variant. The triangle indicates the migration distance of well-formed monomeric structures.

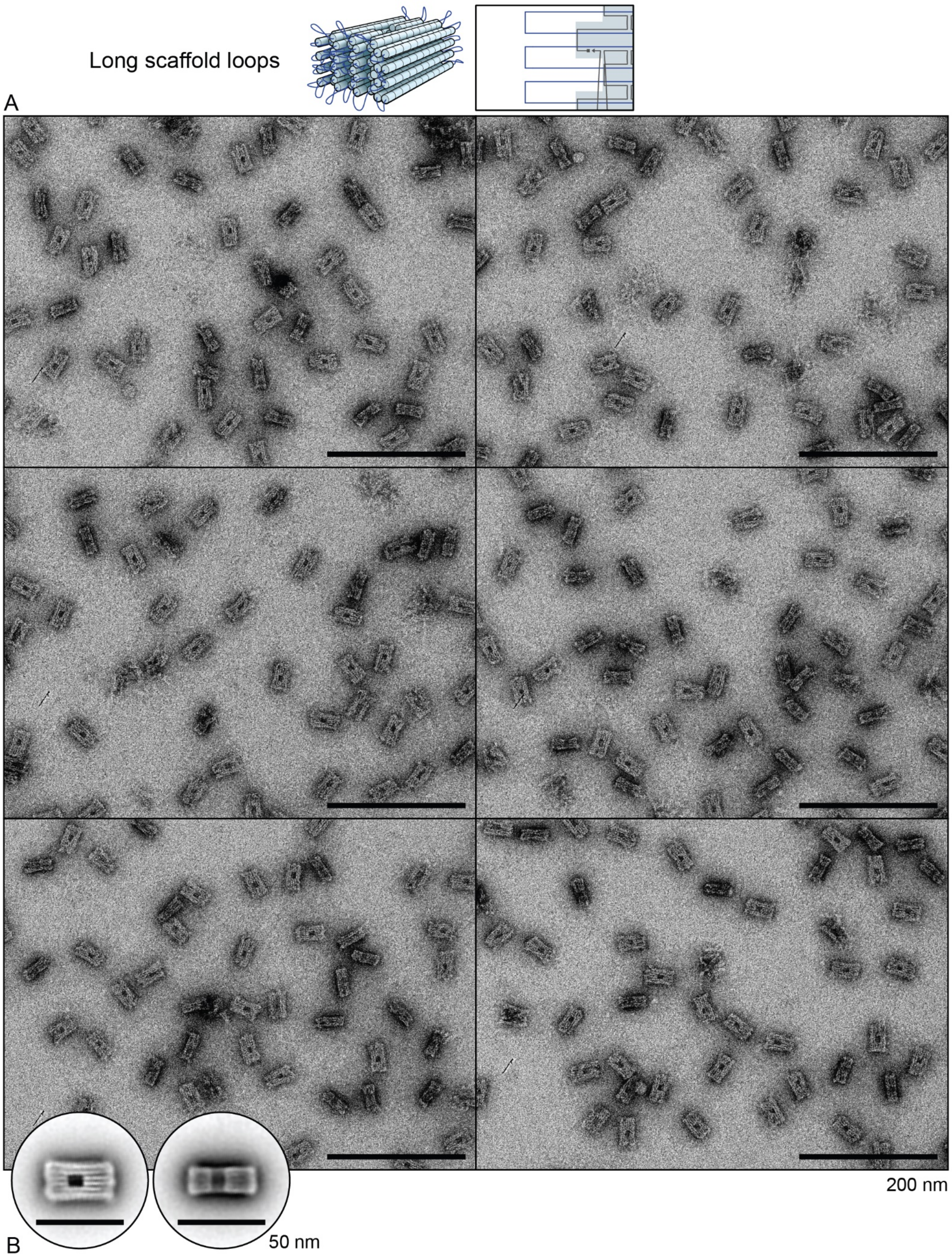


Supplementary figure 14 – Transmission electron micrographs of long scaffold loop variant. **A)** Representative micrographs. **B)** Two-dimensional particle averages.

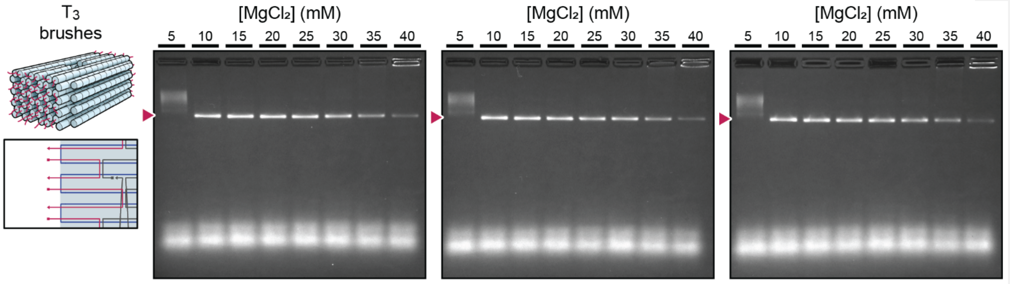


Supplementary figure 15 – Agarose gel electrophoresis results (3 replicates) for $T_3$ brush variant. The triangle indicates the migration distance of well-formed monomeric structures.

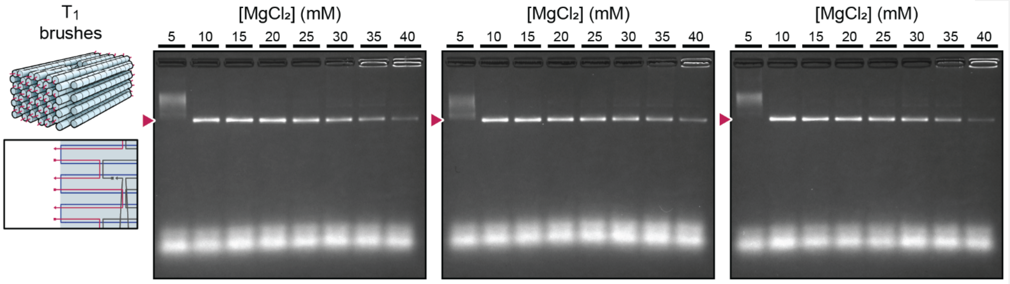


Supplementary figure 16 – Agarose gel electrophoresis results (3 replicates) for $T_1$ brush variant. The triangle indicates the migration distance of well-formed monomeric structures.

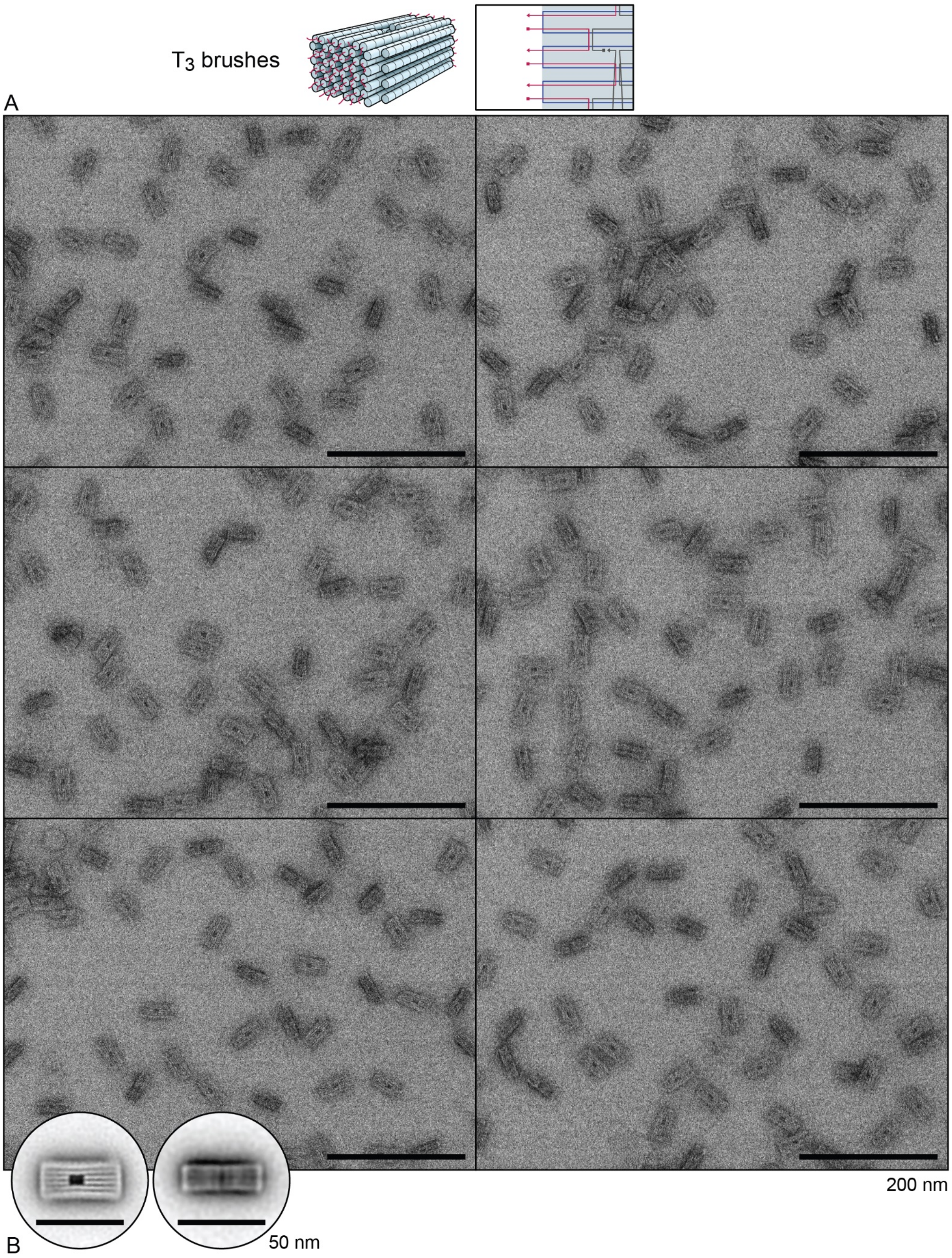


Supplementary figure 17 – Transmission electron micrographs of $T_3$ brush variant. **A)** Representative micrographs. **B)** Two-dimensional particle averages.

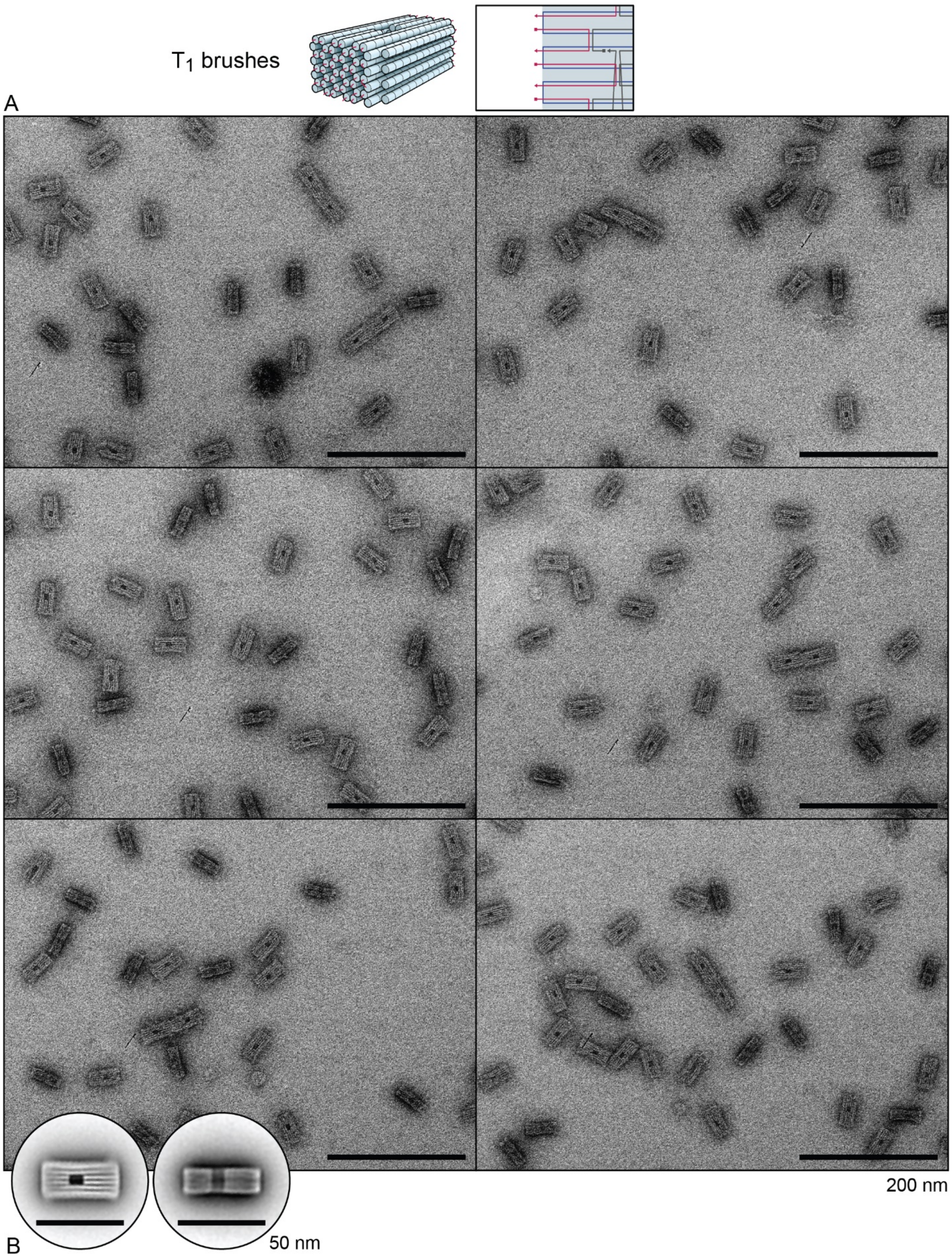


Supplementary figure 18 – Transmission electron micrographs of $T_1$ brush variant. **A)** Representative micrographs. **B)** Two-dimensional particle averages.

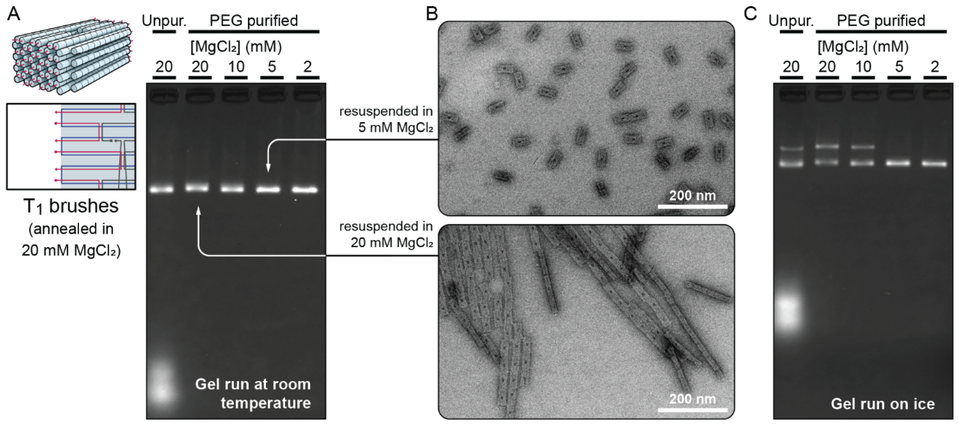


Supplementary figure 19 – Discrepancies between methods when analysing the aggregation of the $T_1$ variant. **A)** Agarose gel electrophoresis results of the $T_1$ variant unpurified, and PEG purified with resuspension in buffer with different concentrations of $MgCl_2$. This gel was set up using initially-refrigerated buffer (4 °C), but then electrophoresed for 3 hours at room temperature (not in an ice bath). **B)** Representative electron micrographs of the $T_1$ variant after PEG purification and resuspension in buffer with 5 mM $MgCl_2$ (top) and 20 mM $MgCl_2$ (bottom). **C)** Agarose gel electrophoresis of the same samples as in A) but electrophoresed for 3 hours in an ice bath.

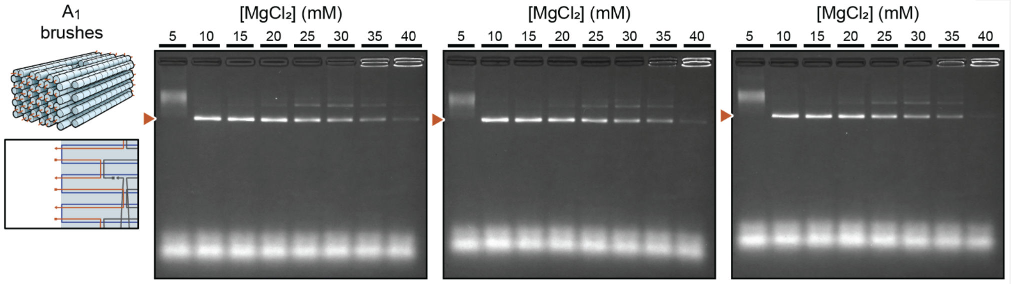


Supplementary figure 20 – Agarose gel electrophoresis results (3 replicates) for $A_1$ brush variant. The triangle indicates the migration distance of well-formed monomeric structures.

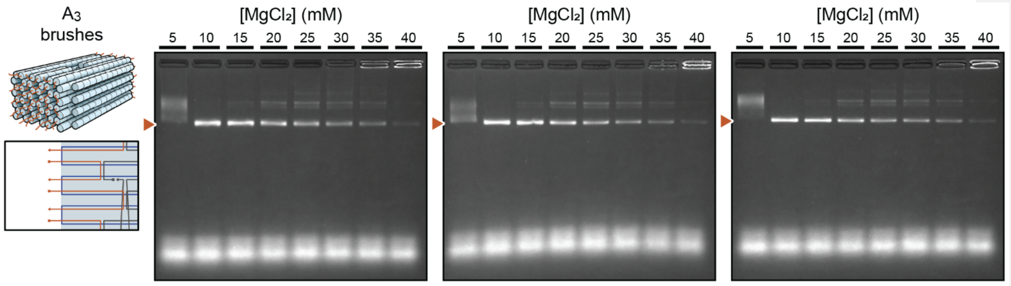


Supplementary figure 21 – Agarose gel electrophoresis results (3 replicates) for $A_3$ brush variant. The triangle indicates the migration distance of well-formed monomeric structures.

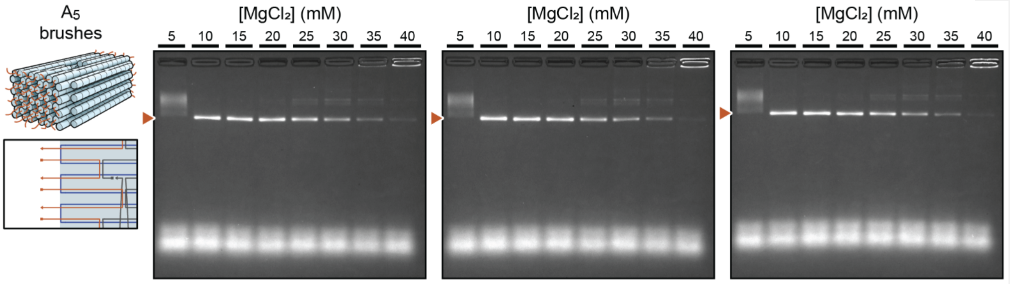


Supplementary figure 22 – Agarose gel electrophoresis results (3 replicates) for $A_5$ brush variant. The triangle indicates the migration distance of well-formed monomeric structures.

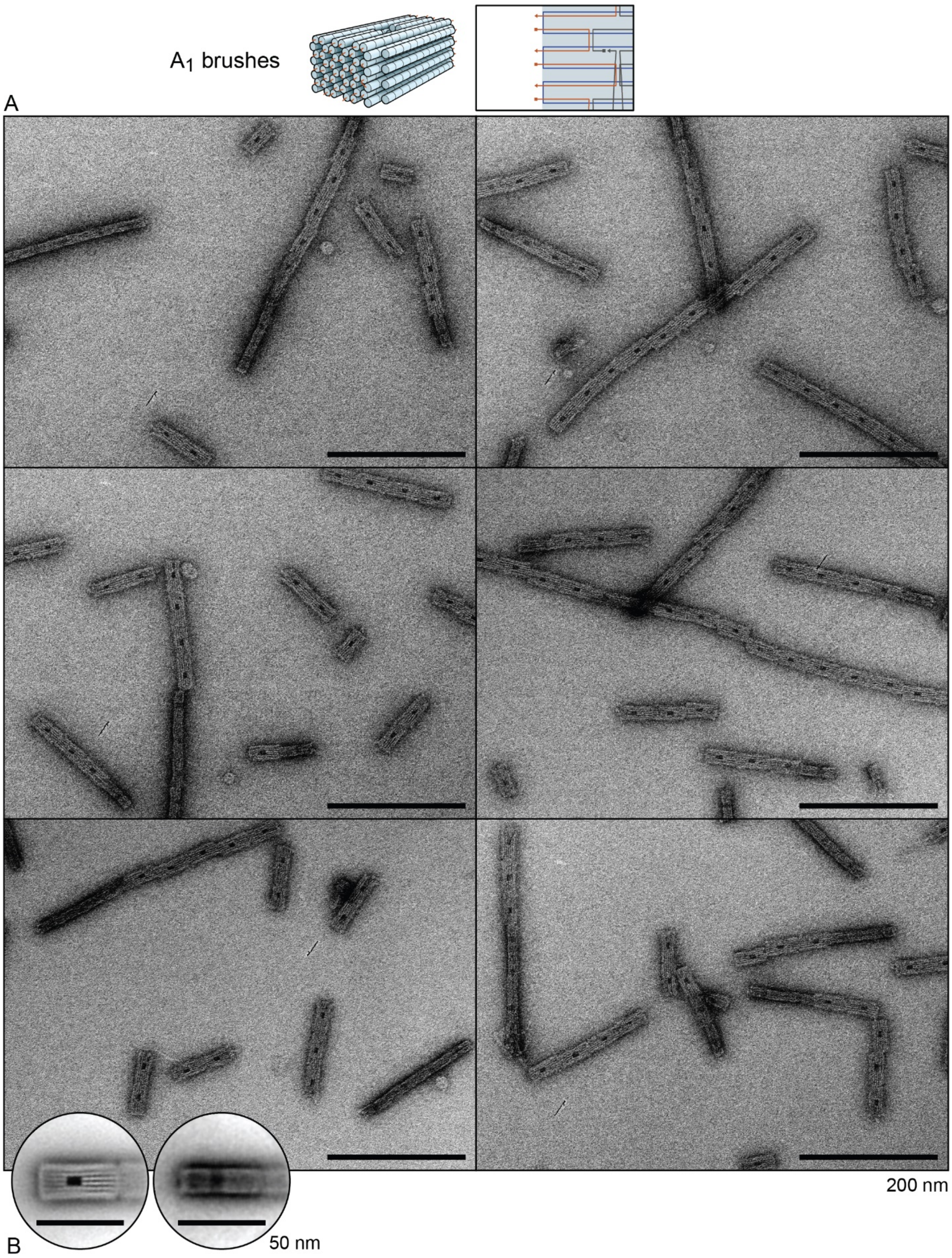


Supplementary figure 23 – Transmission electron micrographs of $A_1$ brush variant. **A)** Representative micrographs. **B)** Two-dimensional particle averages.

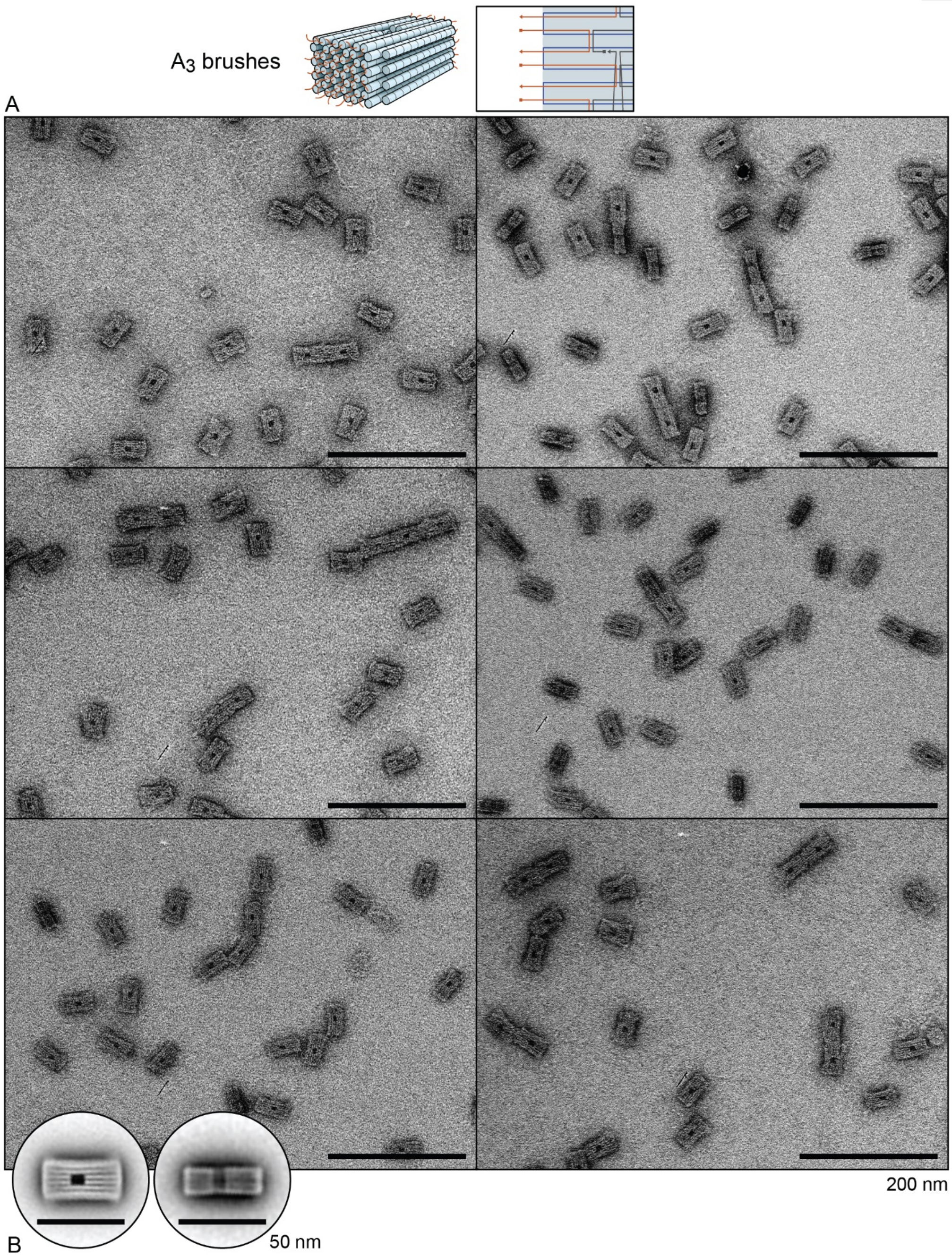


Supplementary figure 24 – Transmission electron micrographs of $A_3$ brush variant. **A)** Representative micrographs. **B)** Two-dimensional particle averages.

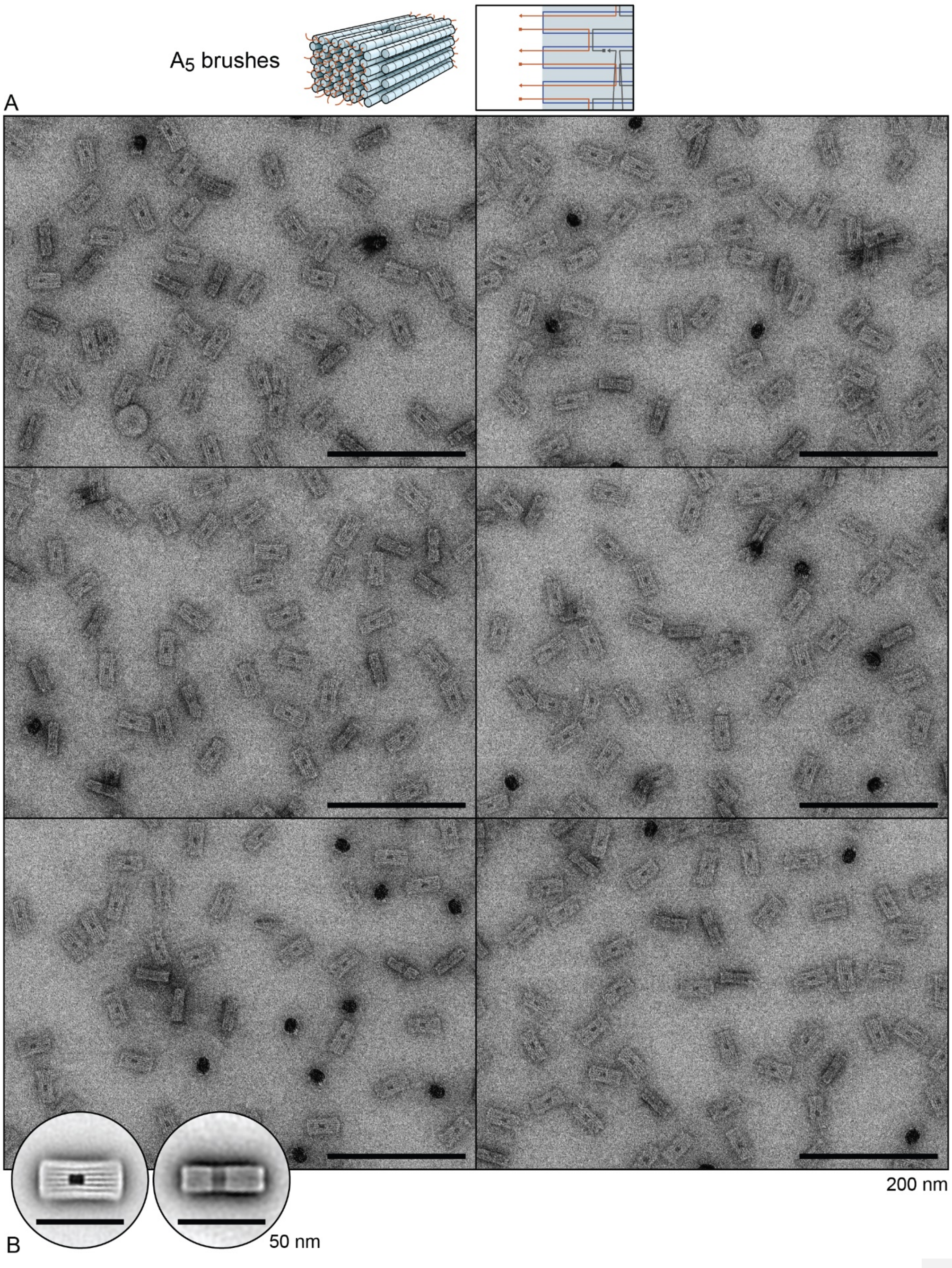


Supplementary figure 25 – Transmission electron micrographs of $A_5$ brush variant. **A)** Representative micrographs. **B)** Two-dimensional particle averages.

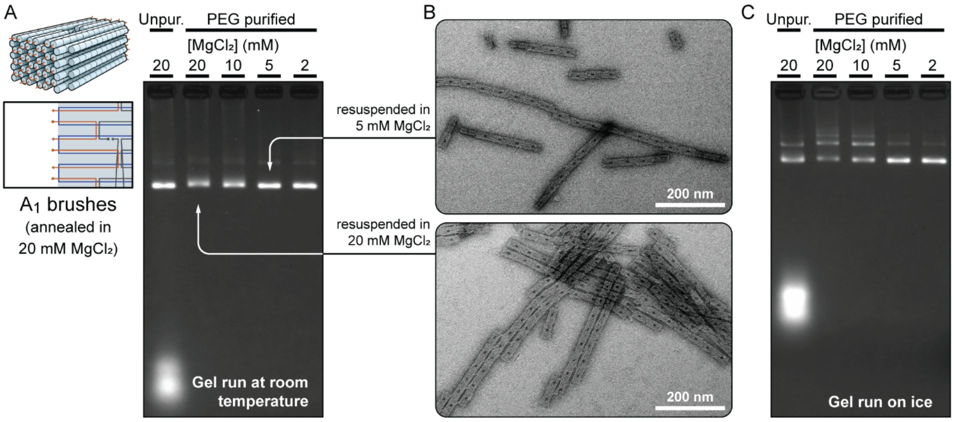


Supplementary figure 26 – Discrepancies between methods when analysing the aggregation of the $A_1$ variant. **A)** Agarose gel electrophoresis results of the $A_1$ variant unpurified, and PEG purified with resuspension in buffer with different concentrations of $MgCl_2$. This gel was set up using initially-refrigerated buffer (4 °C), but then electrophoresed for 3 hours at room temperature (not in an ice bath). **B)** Representative electron micrographs of the $A_1$ variant after PEG purification and resuspension in buffer with 5 mM $MgCl_2$ (top) and 20 mM $MgCl_2$ (bottom). **C)** Agarose gel electrophoresis of the same samples as in A) but electrophoresed for 3 hours in an ice bath.

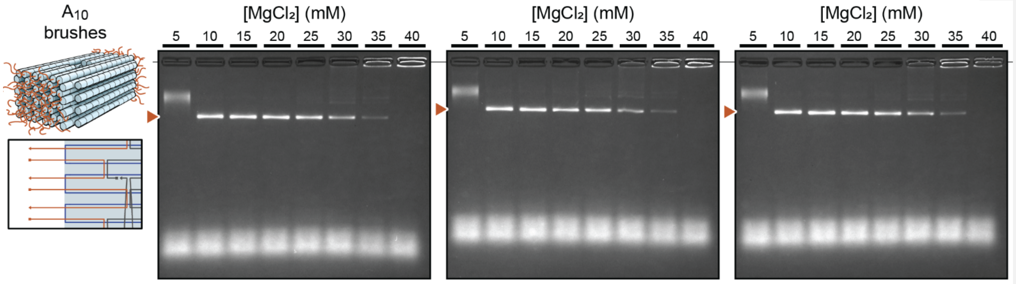


Supplementary figure 27 – Agarose gel electrophoresis results (3 replicates) for $A_{10}$ brush variant. The triangle indicates the migration distance of well-formed monomeric structures.

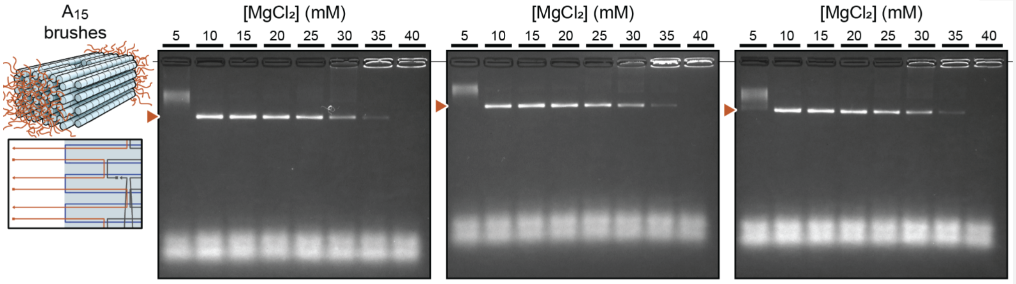


Supplementary figure 28 – Agarose gel electrophoresis results (3 replicates) for $A_{15}$ brush variant. The triangle indicates the migration distance of well-formed monomeric structures.

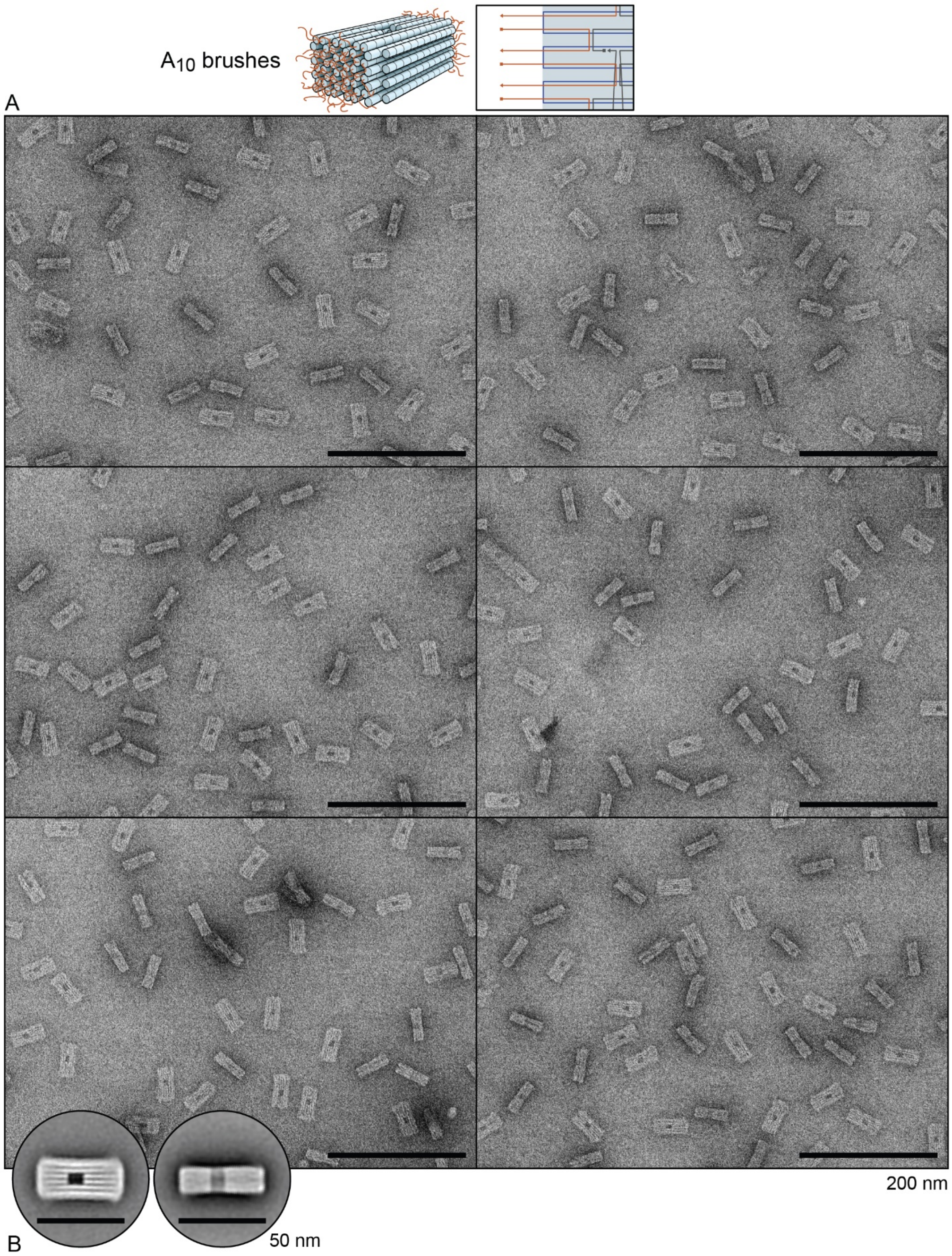


Supplementary figure 29 – Transmission electron micrographs of A$_{10}$ brush variant. **A)** Representative micrographs. **B)** Two-dimensional particle averages.

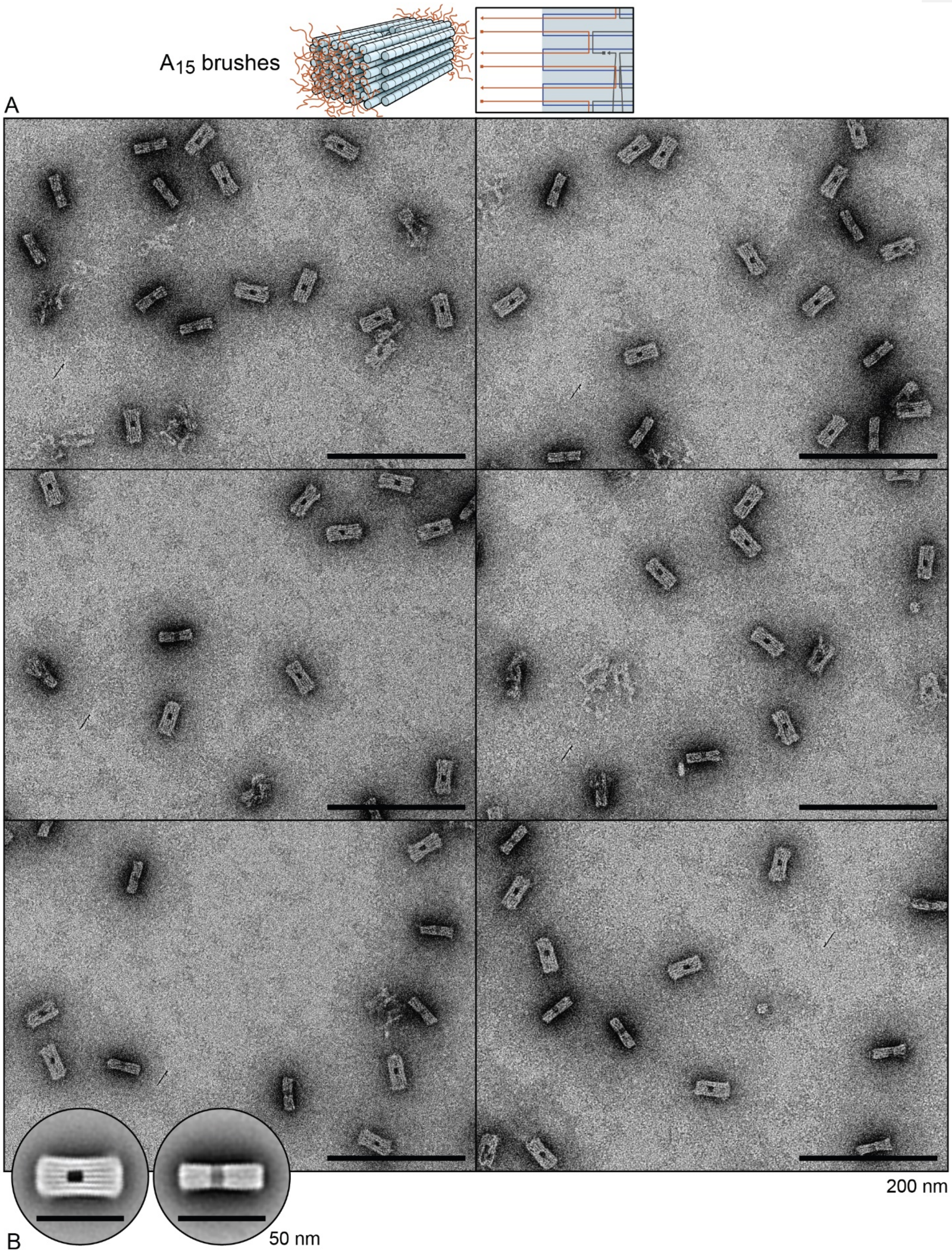


Supplementary figure 30 – Transmission electron micrographs of $A_{15}$ brush variant. **A)** Representative micrographs. **B)** Two-dimensional particle averages.

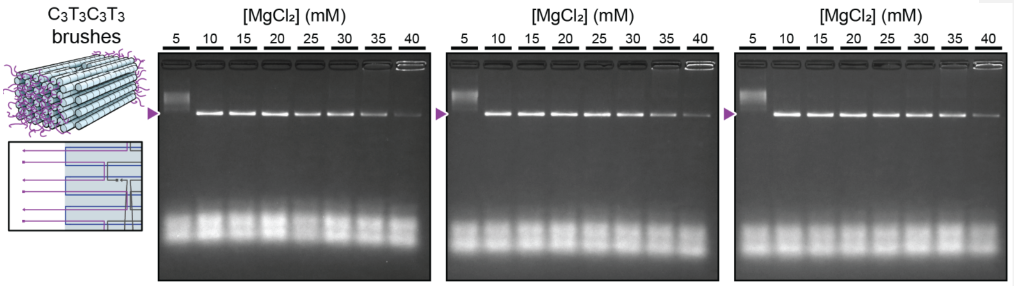


Supplementary figure 31 – Agarose gel electrophoresis results (3 replicates) for $C_3T_3C_3T_3$ brush variant. The triangle indicates the migration distance of well-formed monomeric structures.

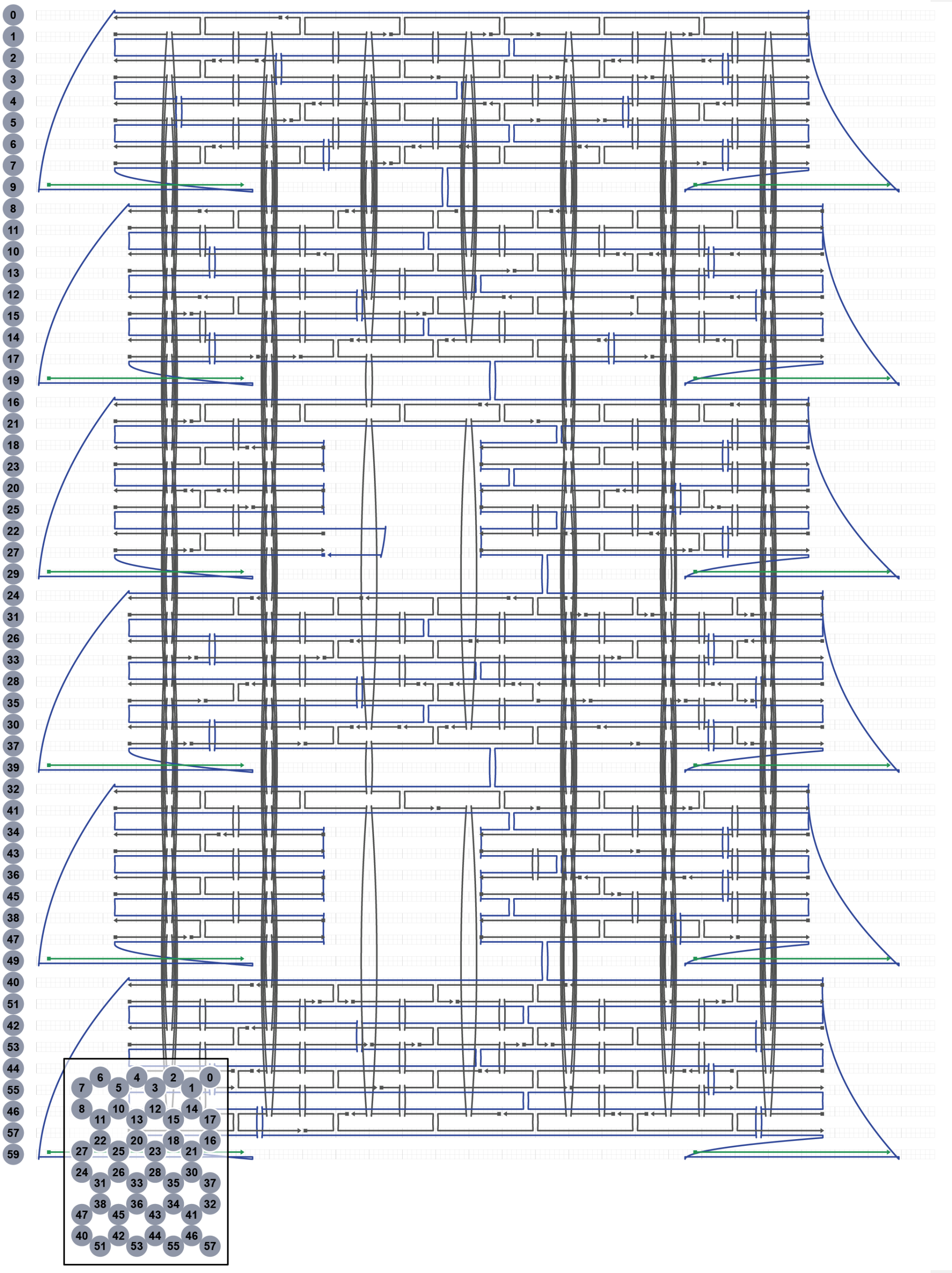

Supplementary figure 32 – Strand schematic layout of orthogonal end-capped variant.

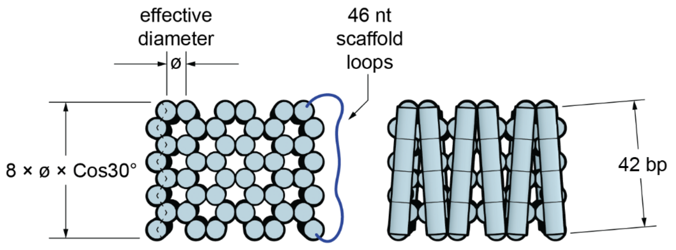


Supplementary figure 33 – Scaffold loop length calculation for orthogonal end-caps. Effective diameter ø has been previously determined to be approximately 2.1 - 2.4 nm, depending on the origami structure[1]. This means that the distance that the scaffold loop (and hence the end-cap helix) must span for each column is between 14.5 and 16.6 nm. We used 42 bp end-caps, (which we estimate to be approximately 14.2 nm when hybridized) with 2 nt unpaired spacers on each side.

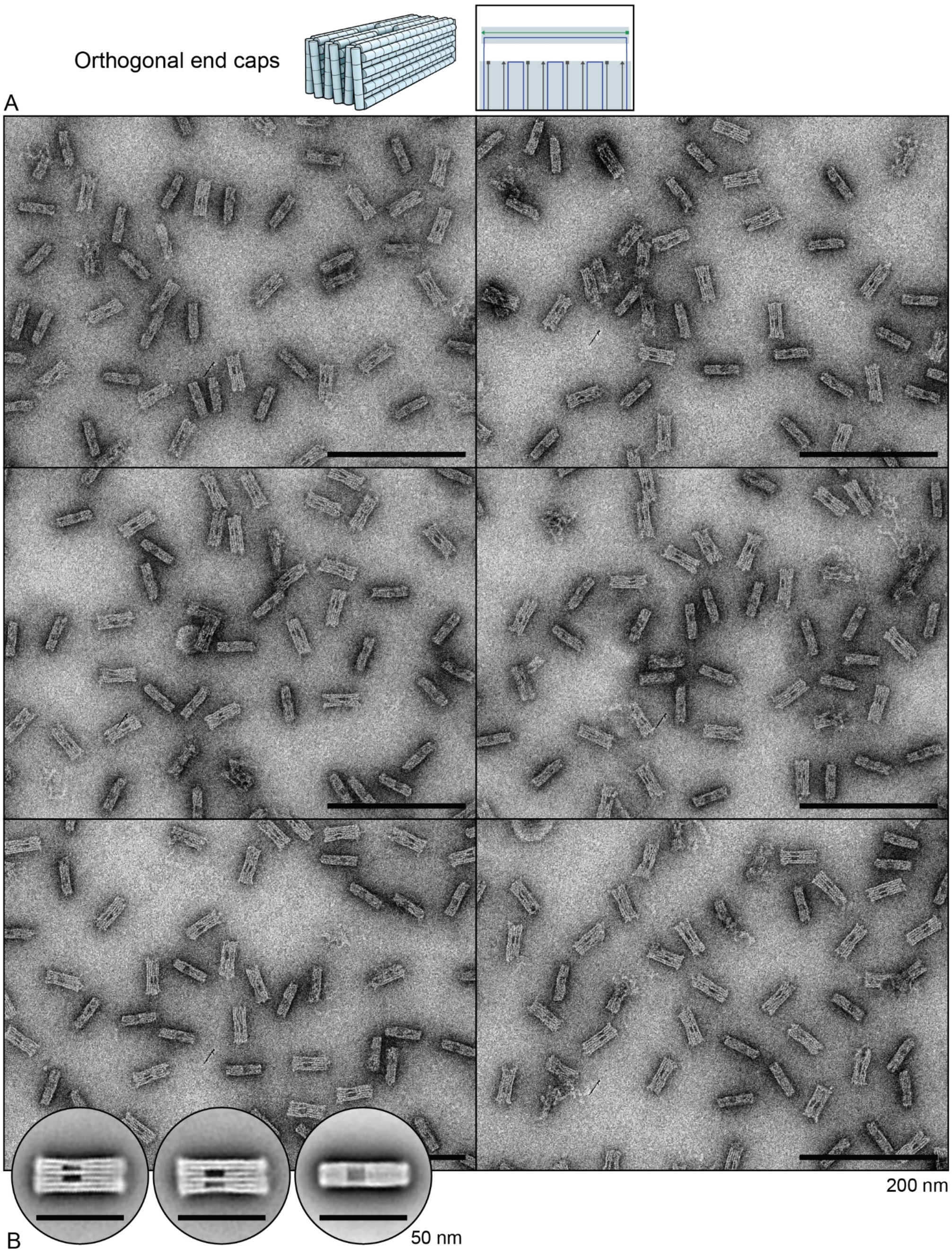


Supplementary figure 34 – Transmission electron micrographs of orthogonal end-capped variant. **A)** Representative micrographs. **B)** Two-dimensional particle averages.

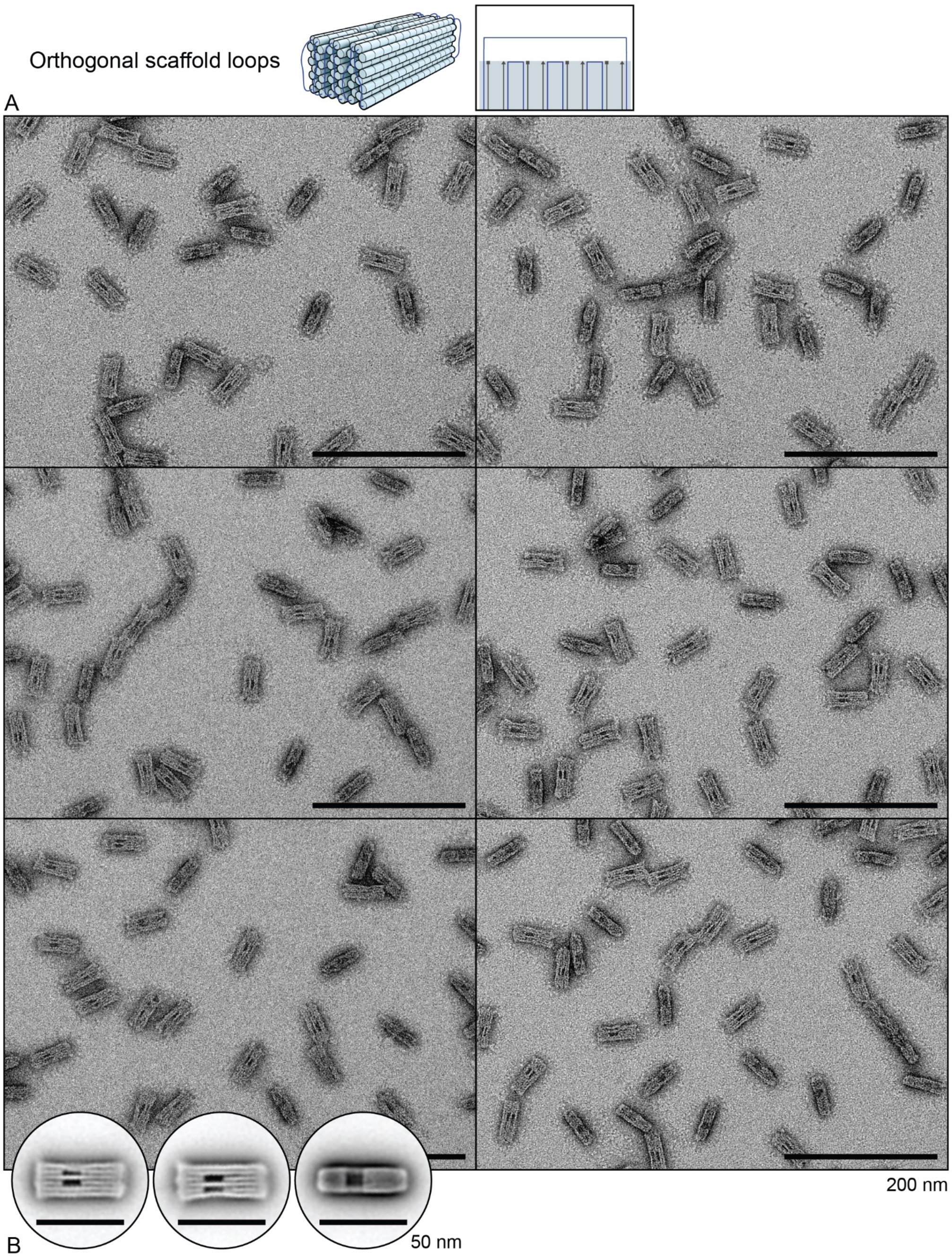


Supplementary figure 35 – Transmission electron micrographs of orthogonal scaffold loop variant. **A)** Representative micrographs. **B)** Two-dimensional particle averages.

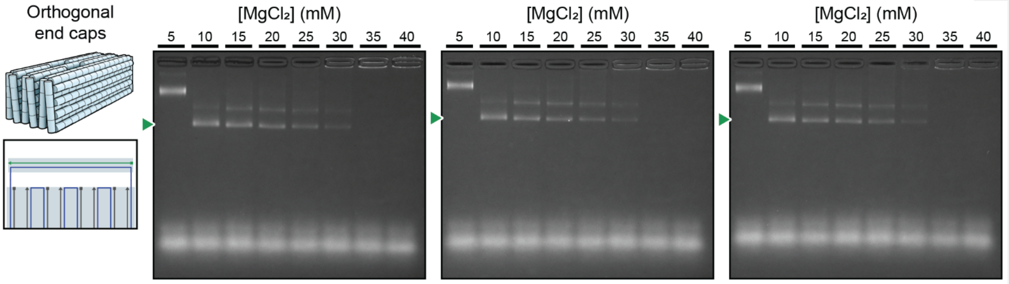


Supplementary figure 36 – Agarose gel electrophoresis results (3 replicates) for the orthogonal end-cap variant. The triangle indicates the migration distance of well-formed monomeric structures.

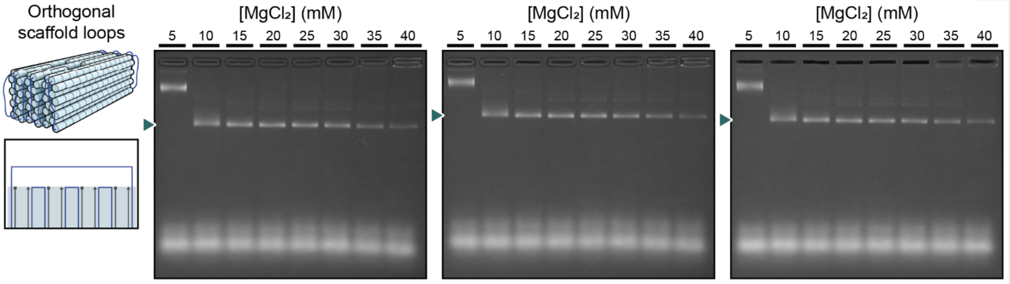


Supplementary figure 37 – Agarose gel electrophoresis results (3 replicates) for the orthogonal scaffold loop variant. The triangle indicates the migration distance of well-formed monomeric structures.

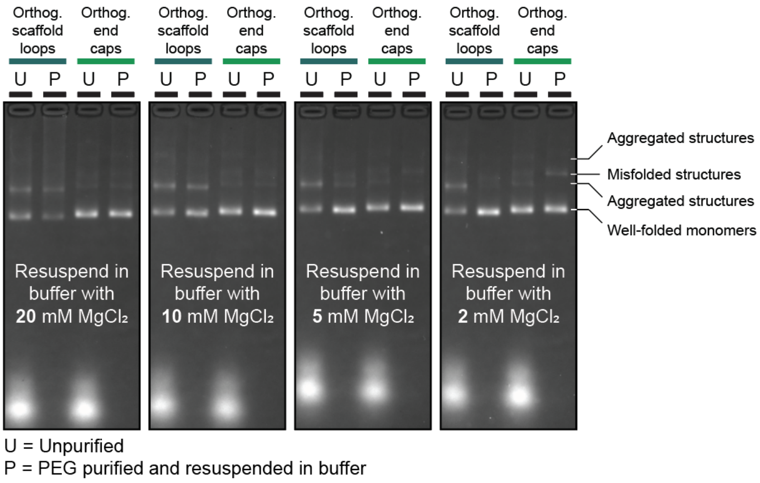


Supplementary figure 38 - Agarose gel results for orthogonal scaffold loop and end-capped DNA origami variants before and after purification and buffer exchange into lower $MgCl_2$.

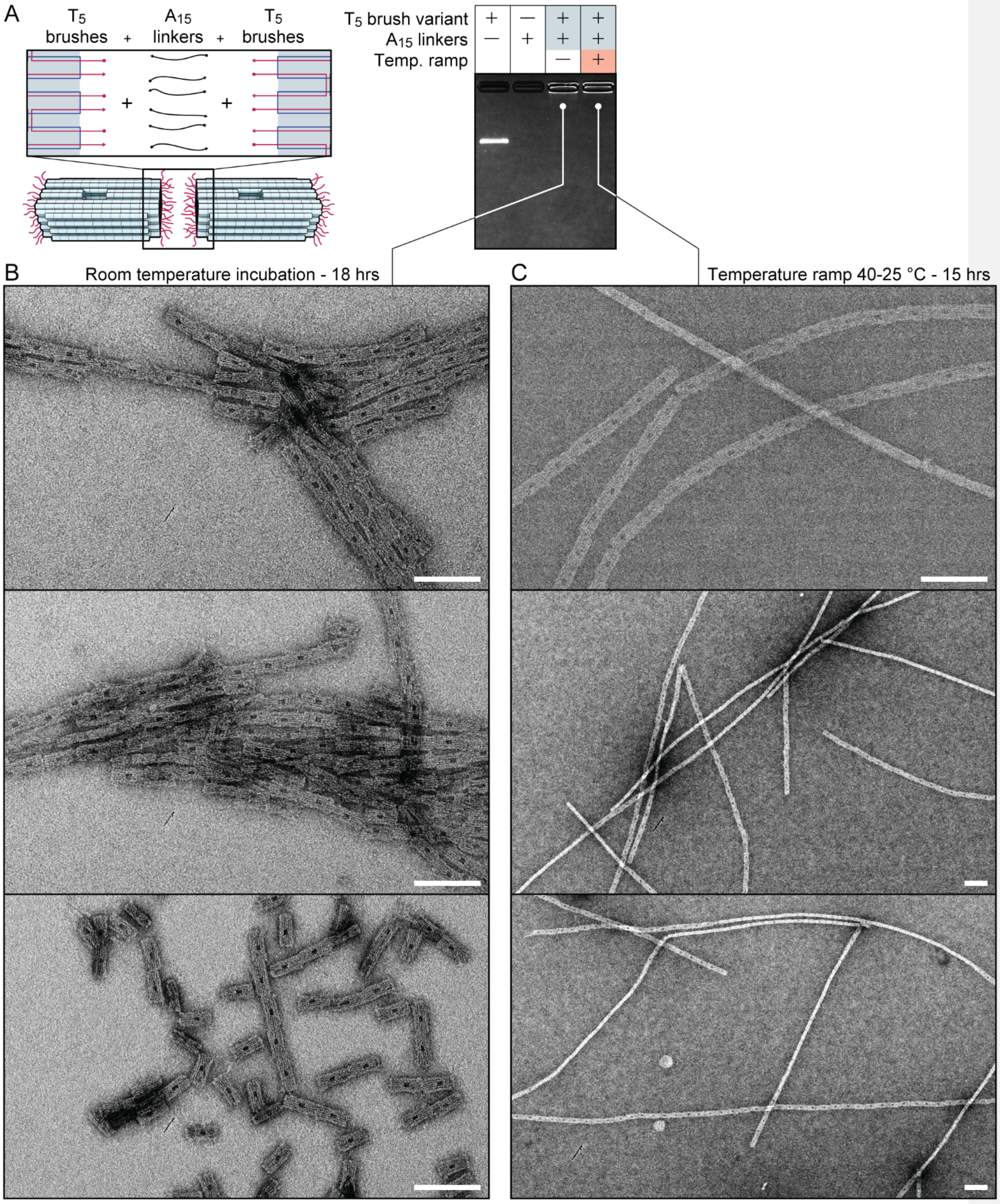


Supplementary figure 39 – Multimeric assembly of the $T_5$ variant with $A_{15}$ linker strands. **A)** Schematic and AGE results showing aggregation in the loading wells. **B)** Transmission electron micrographs of the mixtures incubated at room temperature for 18 hours prior to TEM grid staining. **C)** Transmission electron micrographs of the mixtures that were temperature ramped from 40 °C to 25 °C over 15 hours prior to TEM grid staining. All scale bars are 100 nm.

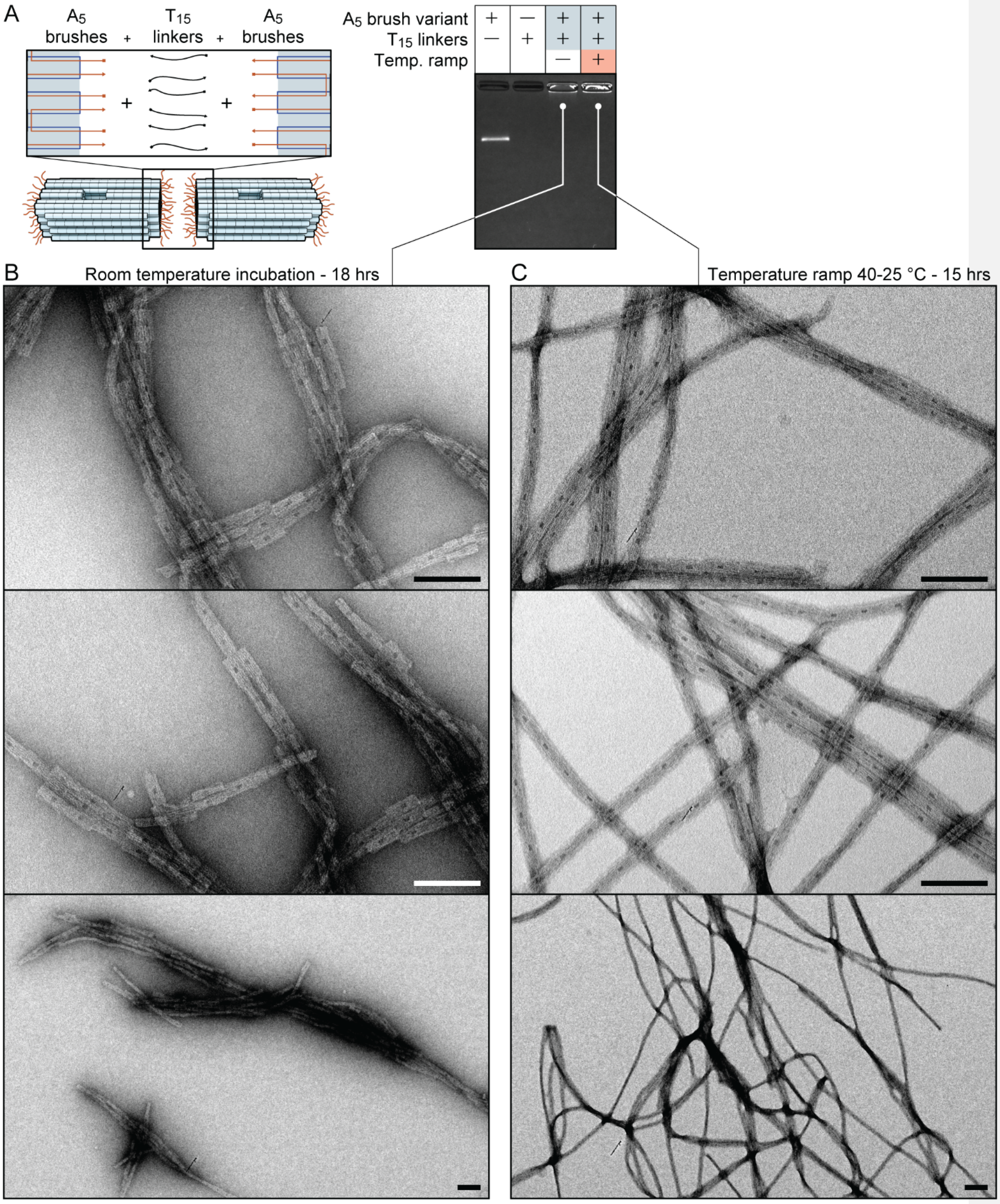


Supplementary figure 40 – Multimeric assembly of the $A_5$ variant with $T_{15}$ linker strands. **A)** Schematic and AGE results showing aggregation in the loading wells. **B)** Transmission electron micrographs of the mixtures incubated at room temperature for 18 hours prior to TEM grid staining. **C)** Transmission electron micrographs of the mixtures that were temperature ramped from 40 °C to 25 °C over 15 hours prior to TEM grid staining. All scale bars are 100 nm.

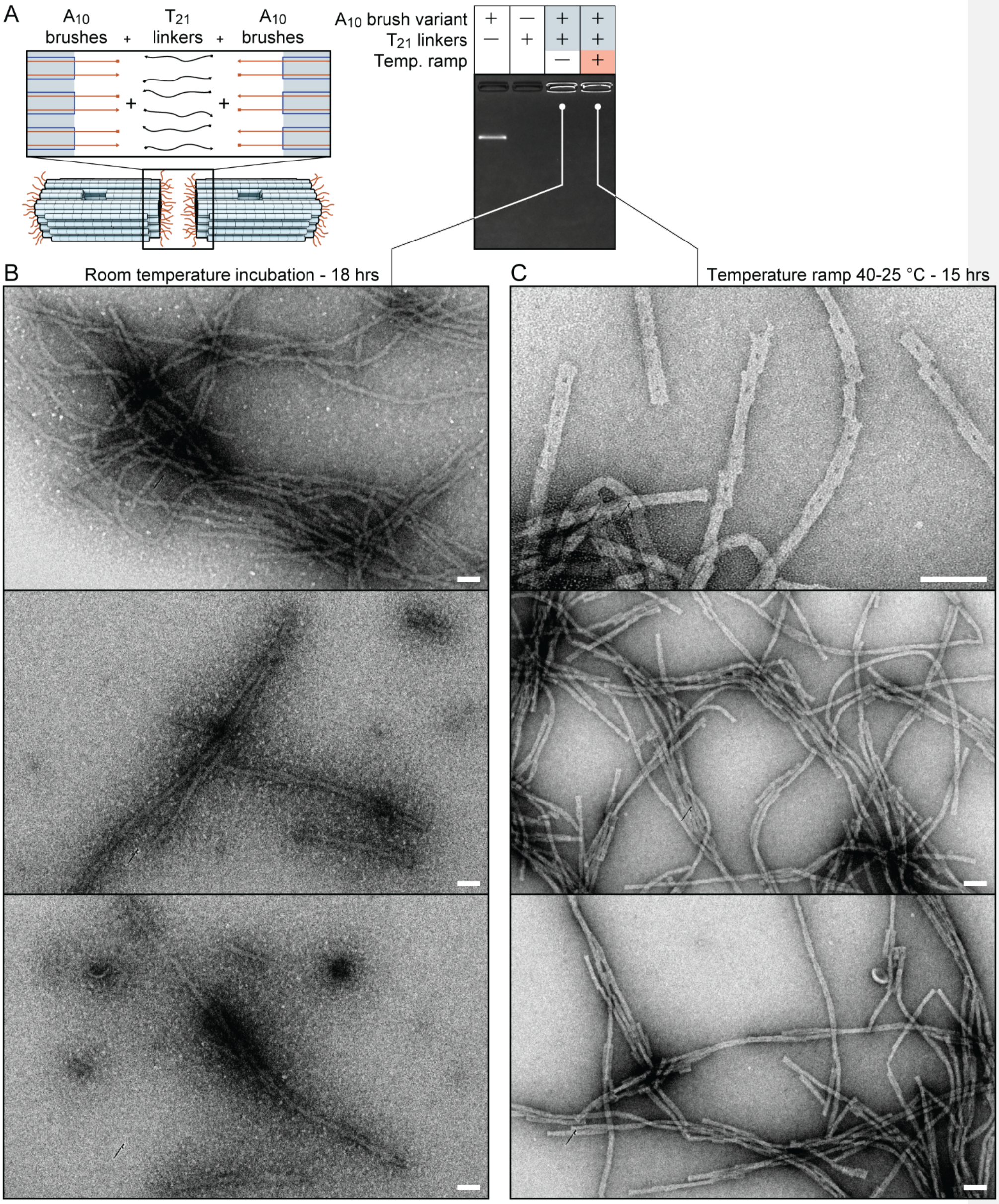


Supplementary figure 41 – Multimeric assembly of the $A_{10}$ variant with $T_{21}$ linker strands. **A)** Schematic and AGE results showing aggregation in the loading wells. **B)** Transmission electron micrographs of the mixtures incubated at room temperature for 18 hours prior to TEM grid staining. **C)** Transmission electron micrographs of the mixtures that were temperature ramped from 40 °C to 25 °C over 15 hours prior to TEM grid staining. All scale bars are 100 nm.

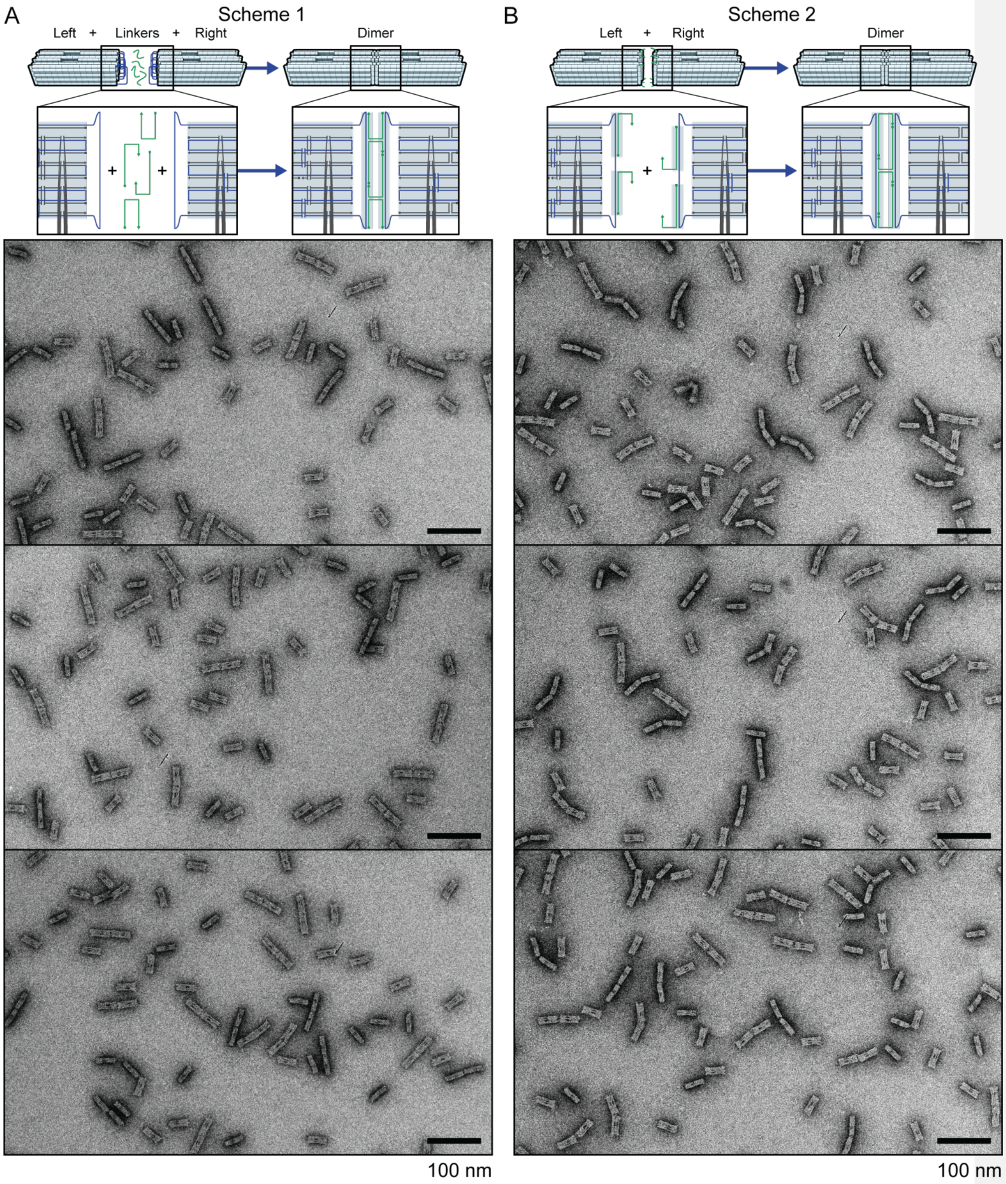


Supplementary figure 42 – Schematics and TEM micrographs of OEC dimer assembly schemes. **A)** Top: Explanation of scheme 1, where linker strands are added after the left and right subunits are synthesized, purified and mixed. Bottom: Exemplary micrographs. **B)** Top: Explanation of scheme 2, where left and right subunits are synthesized, purified and mixed. Bottom: Exemplary micrographs.

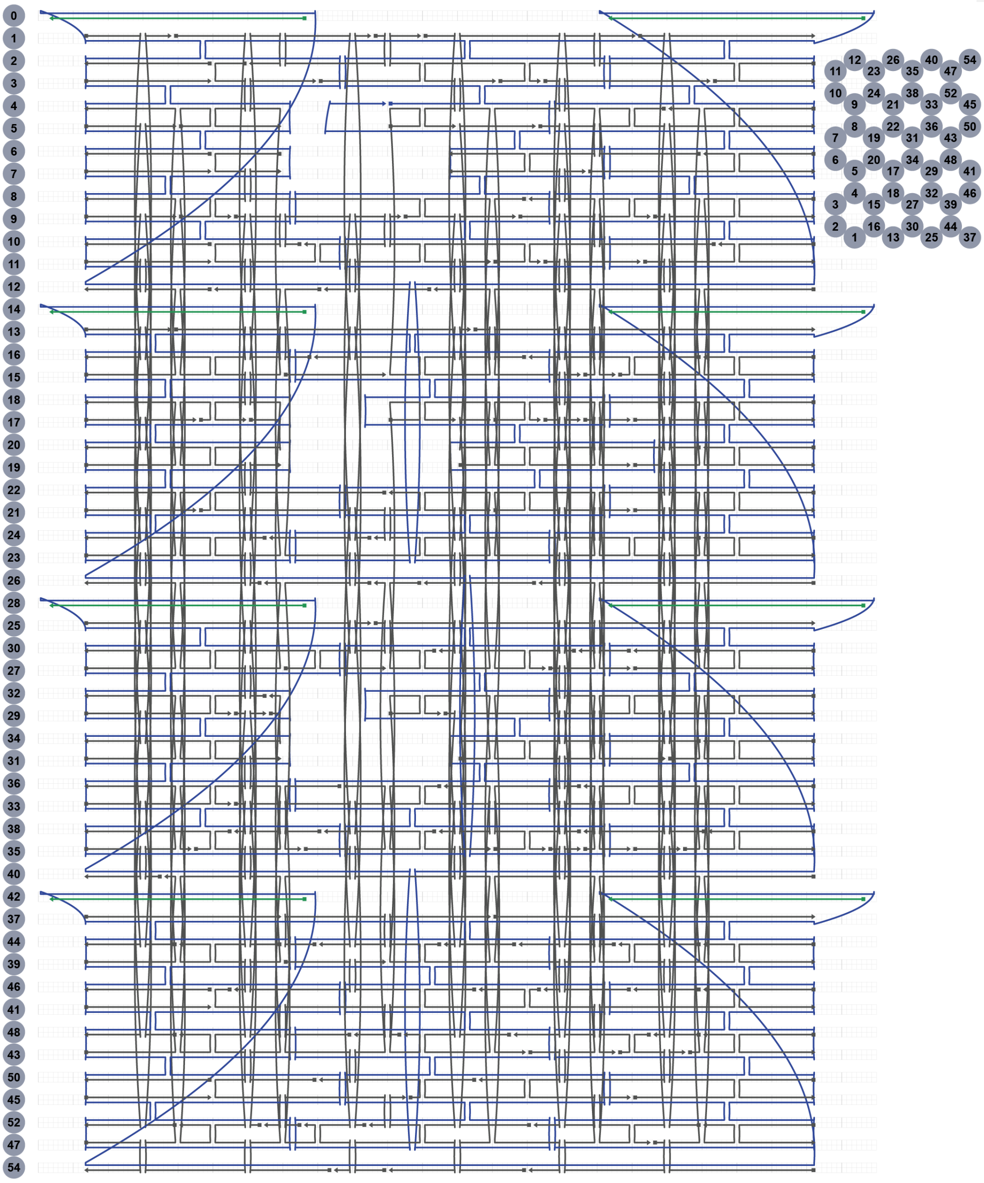


Supplementary figure 43 – Strand schematic layout of orthogonal end-capped variant (horizontal version 1)

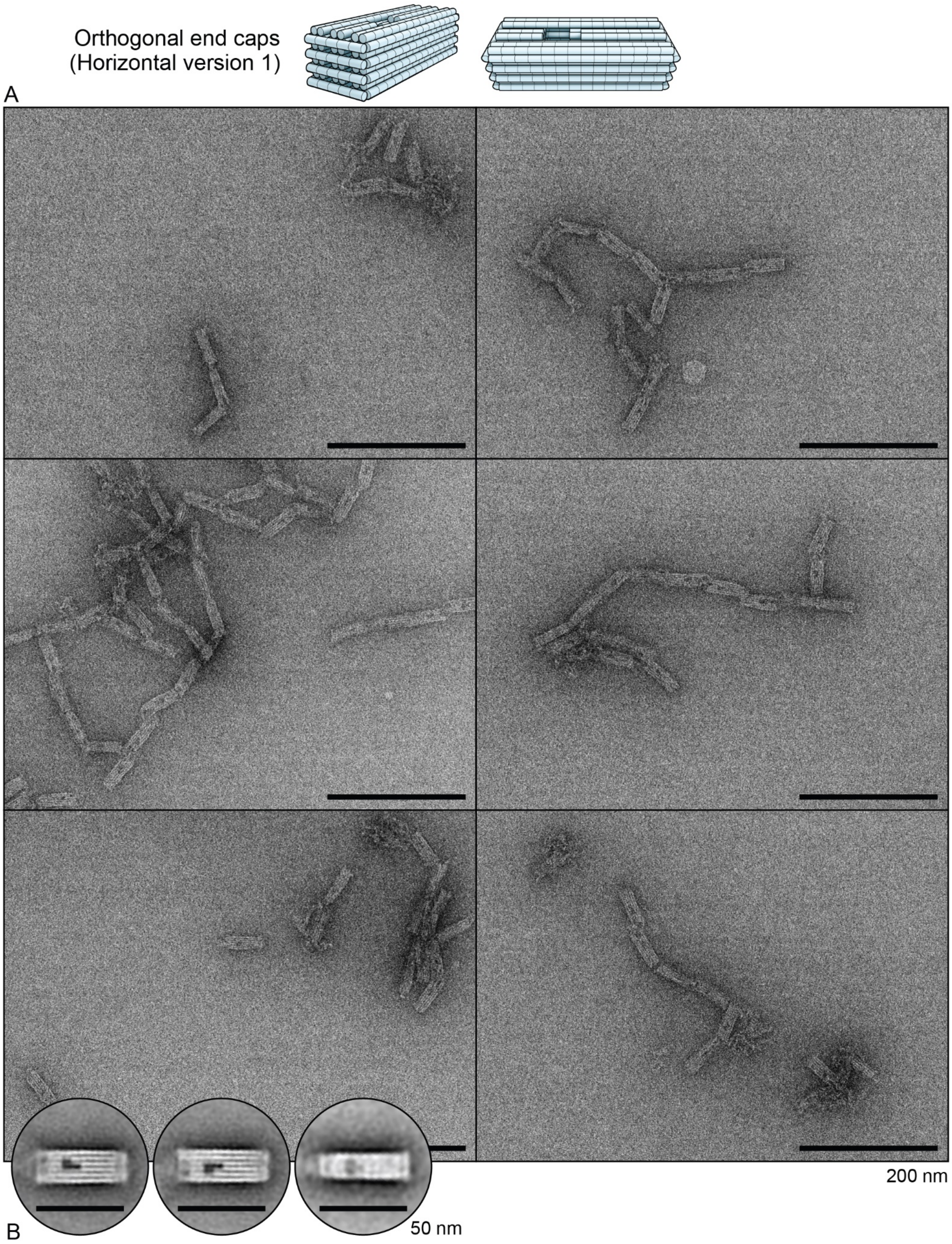


Supplementary figure 44 – Transmission electron micrographs of orthogonal end-capped variant (horizontal version 1). **A)** Representative micrographs. **B)** Two-dimensional particle averages.

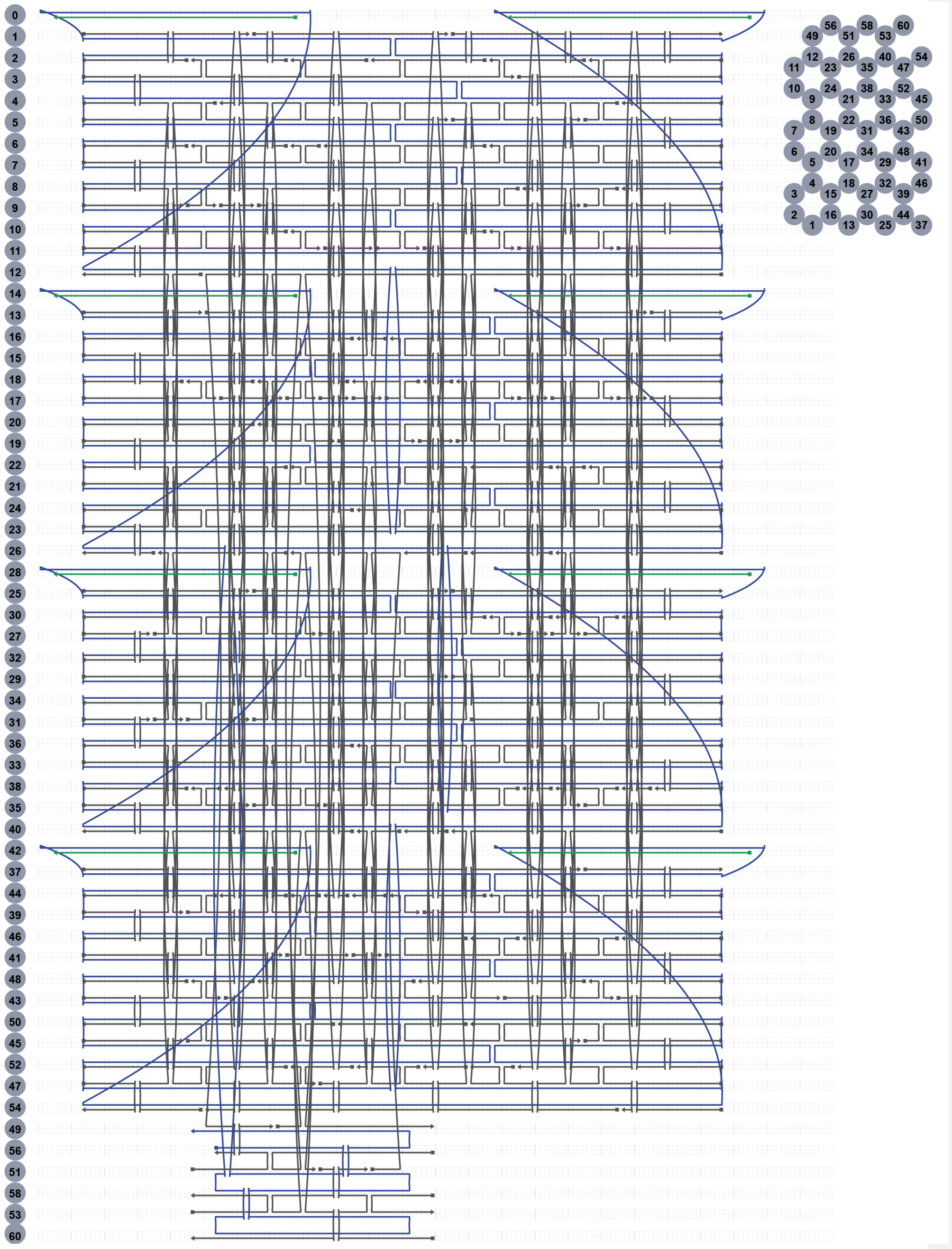


Supplementary figure 45 – Strand schematic layout of orthogonal end-capped variant (horizontal version 2)

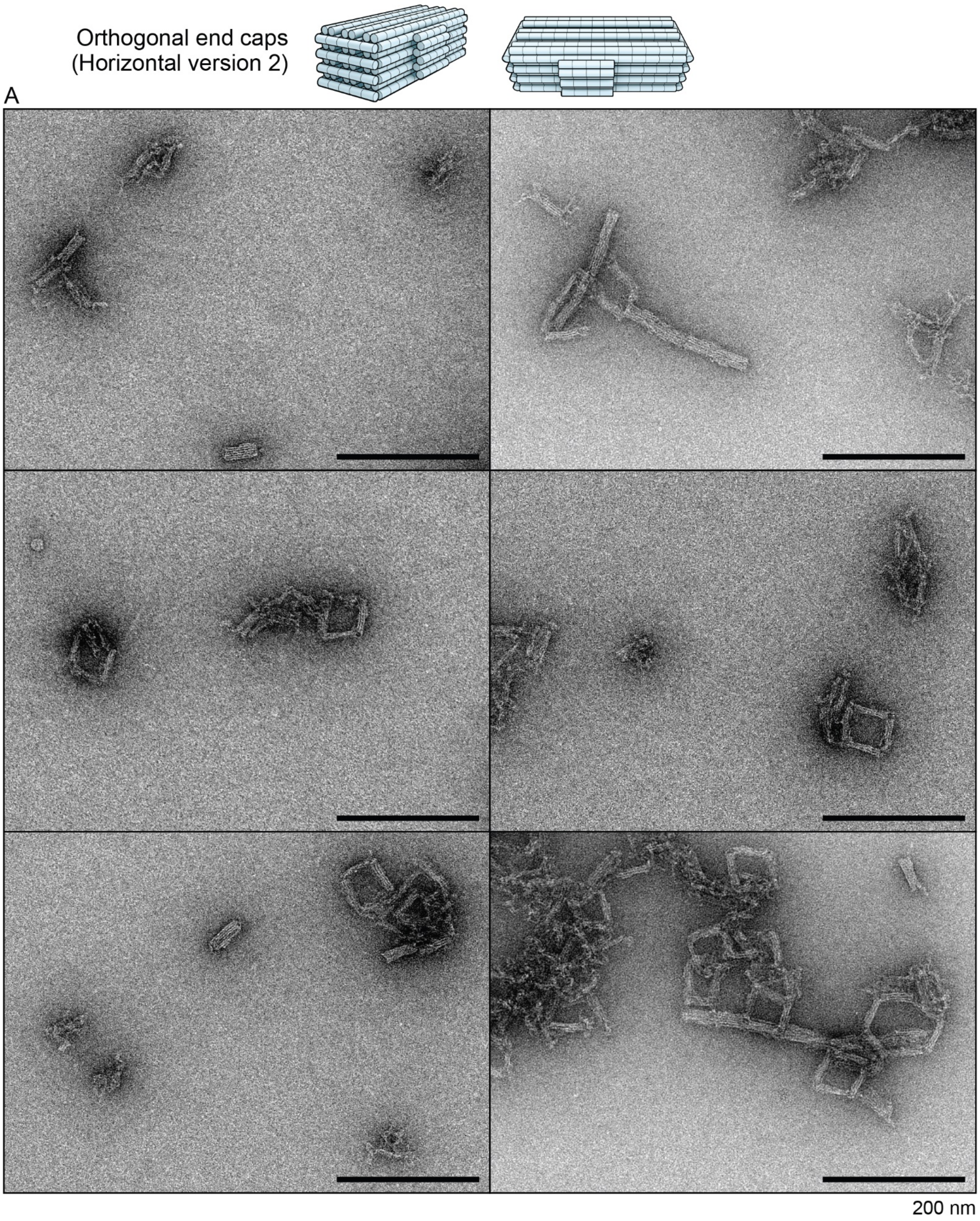


Supplementary figure 46 – Transmission electron micrographs of orthogonal end-capped variant (horizontal version 2). **A)** Representative micrographs.
